# Viral genomic features predict *orthopoxvirus* reservoir hosts

**DOI:** 10.1101/2023.10.26.564211

**Authors:** Katie K. Tseng, Heather Koehler, Daniel J. Becker, Rory Gibb, Colin J. Carlson, Maria del Pilar Fernandez, Stephanie N. Seifert

**Affiliations:** Paul G. Allen School for Global Health, Washington State University, Pullman, Washington, United States of America; School of Molecular Biosciences, Washington State University, Pullman, Washington, United States of America; School of Biological Sciences, University of Oklahoma, Norman, Oklahoma, United States of America; Centre for Biodiversity and Environment Research, Department of Genetics, Evolution and Environment, University College London, London, United Kingdom; People & Nature Lab, UCL East, University College London, Stratford, London, United Kingdom; Center for Global Health Science and Security, Georgetown University, Washington, DC, United States of America

**Keywords:** Infectious diseases, prediction, machine learning, boosted regression trees, orthopoxviruses, Mpox

## Abstract

Orthopoxviruses (OPVs), including the causative agents of smallpox and mpox have led to devastating outbreaks in human populations worldwide. However, the discontinuation of smallpox vaccination, which also provides cross-protection against related OPVs, has diminished global immunity to OPVs more broadly. We apply machine learning models incorporating both host ecological and viral genomic features to predict likely reservoirs of OPVs. We demonstrate that incorporating viral genomic features in addition to host ecological traits enhanced the accuracy of potential OPV host predictions, highlighting the importance of host-virus molecular interactions in predicting potential host species. We identify hotspots for geographic regions rich with potential OPV hosts in parts of southeast Asia, equatorial Africa, and the Amazon, revealing high overlap between regions predicted to have a high number of potential OPV host species and those with the lowest smallpox vaccination coverage, indicating a heightened risk for the emergence or establishment of zoonotic OPVs. Our findings can be used to target wildlife surveillance, particularly related to concerns about mpox establishment beyond its historical range.

## Introduction

*Variola virus*, the smallpox-causing agent belonging to the *Orthopoxvirus* genus (OPV), has left an indelible mark on human history. This exceptionally virulent disease has triggered some of the most catastrophic outbreaks in human memory, leading to significant morbidity and mortality on a global scale. However, it also catalyzed progress in therapeutic intervention, inspiring the development of the first and highly effective vaccine. This vaccine leveraged cross-protective immunity from a closely related but less virulent OPV, ultimately leading to the successful eradication of smallpox[1]. In 1980, smallpox became the first human disease to be eradicated as a result of a global, coordinated vaccination effort --- and only made possible by a lack of suitable animal reservoirs to maintain variola virus outside of human populations[2]. After the eradication of smallpox, vaccinations were largely discontinued, resulting in a worldwide reduction in immunological memory against OPVs[3]. Nevertheless, the OPV genus is a diverse group, with many members still circulating in animal reservoirs, facilitating periodic emergence in humans and complicating intervention efforts.

Orthopoxviruses are notable for their varied mammalian host breadth and pathogenicity, although the full range of circulation in wildlife is unknown [4]. The diversity in host breath has been attributed to variations in the large repertoire of accessory genes found across poxvirus species including many genes associated with the inhibition of host innate immune responses [6–8]. The evolutionary dynamics of OPVs is marked by gains and losses among accessory genes, with genome reduction thought to contribute to host specialization [4]. For example, the modified vaccinia virus Ankara, a third-generation smallpox vaccine, lost considerable genetic information (roughly 15% of the ∼200kb genome), including many genes used by OPVs to regulate the mammalian host environment, resulting in a restricted host range [9]. The viral genome of cetaceanpox viruses, a putative sister clade to OPVs, encodes for roughly half the number of proteins found in other poxviruses (7), and these viruses are thought to have highly restricted host ranges. Other OPVs, including mpox virus (formerly known as monkeypox virus) and cowpox virus, are capable of infecting a broad range of hosts, increasing the likelihood of recurrent spillover events into the human population [4]. The unprecedented recent global spread of mpox virus has raised concerns about human-to-animal spillback to susceptible hosts outside its historical range, as has been observed with SARS-CoV-2 in white-tailed deer populations [11]. Furthermore, the recent emergence of the novel borealpox virus (formerly known as alaskapox virus) in humans from an unknown animal reservoir suggests a high likelihood of unidentified OPVs in nature with zoonotic potential [12].

These types of modeling studies may be undermined by their reliance on host ecological and phylogenetic features, and conversely, the lack of predictors that capture host-virus compatibility at molecular scales. For example, several such modeling efforts have indicated a high probability of compatibility between domestic pigs (*Sus scrofa*) and SARS-CoV-2 [16,18,19], which was further supported by *in vitro* susceptibility of pig-derived cell lines [20].

However, *in vivo* challenge studies demonstrated that SARS-CoV-2 fails to infect domestic pigs [21,22], suggesting a post-entry incompatibility that is poorly understood or captured in current predictive models. Similarly, the Egyptian fruit bat, *Rousettus aegyptiacus*, was predicted to be a compatible host for Nipah virus using host trait data [23], but failed to support Nipah virus replication with experimental challenge [24]. Furthermore, viruses evolve over time, changing host breadth or transmission potential, as illustrated by the expanded host range of the SARS-CoV-2 Omicron variant [25]. As the evolution of viruses is shaped by the host environment, from cell entry to evading host innate and adaptive immune response, viral genomes encode valuable information for predicting their host range but have seldom been included in predictive modeling of host-virus associations.

Here, we developed a trait-based approach using boosted regression trees (BRTs), a machine learning algorithm commonly used in ecological and evolutionary research, and integrated both host ecological traits and viral genomic features to predict mammal-OPV associations. To provide a basis for comparison, we constructed traditional host prediction models that used only host traits to predict OPV positivity across mammal genera. These models, trained separately on datasets of candidate host genera with evidence of exposure to OPV species (based on molecular detection of viral genetic material) or likely susceptible to infection (based on isolation of an OPV from a host), assess how different detection methods influence predictions. Building on this framework, we combined PCR and virus isolation data with OPV whole genome sequences to create a comprehensive dataset of host-virus interactions. Using this enriched dataset, we developed a model to predict the probability of a specific link between a single OPV and a single host genus (i.e., *link prediction*) by including both host traits and virus features critical to host-virus interactions on a molecular scale. Unlike a host prediction model, which predicts positivity for a virus or group of viruses, the link prediction model predicts compatible host-virus pairs, an approach that can be applied to optimize targeted sampling and provide quantitative insights into the hypothesized role of viral genomic variation in shaping OPV host range.

## Results

### Host prediction models

Host prediction BRT models aimed at predicting the host range of OPVs using either known host exposure to OPVs (based on PCR) or known susceptible hosts of OPVs (based on virus isolation) had moderate predictive accuracy (Fig 1; S1 Table). Compared to the host exposure model, the susceptible host model had higher overall model accuracy (area under the receiver operating characteristic curve [AUC] = 0.88 vs. 0.86; *t* = -4.38, *p*<0.001), higher specificity (t = - 7.21, *p <* 0.001), and lower sensitivity (*t =* 5.66, *p* < 0.001) (Fig 1; S1 Table). Likewise, the predicted probabilities of OPV positivity were significantly correlated between the two evidence type models (Spearman ρ = 0.535*, p*<0.001), with predictions of the host exposure model and the susceptible host model both exhibiting strong phylogenetic signals (Pagel’s λ = 0.80 and 0.90, respectively). Phylogenetic factorization identified some overlapping taxonomic patterns in the predictions of both host prediction models, including a higher mean probability of genera from the family Felidae (*n* = 17 species) to host OPVs, and a lower mean probability of the orders Lagomorpha and Rodentia (*n =* 514) as well as a subclade of the rodent suborder Hystricomorpha (*n* = 74) to host OPVs (Figs 2 and S1; S2 and S3 Tables).

**Fig 1.**
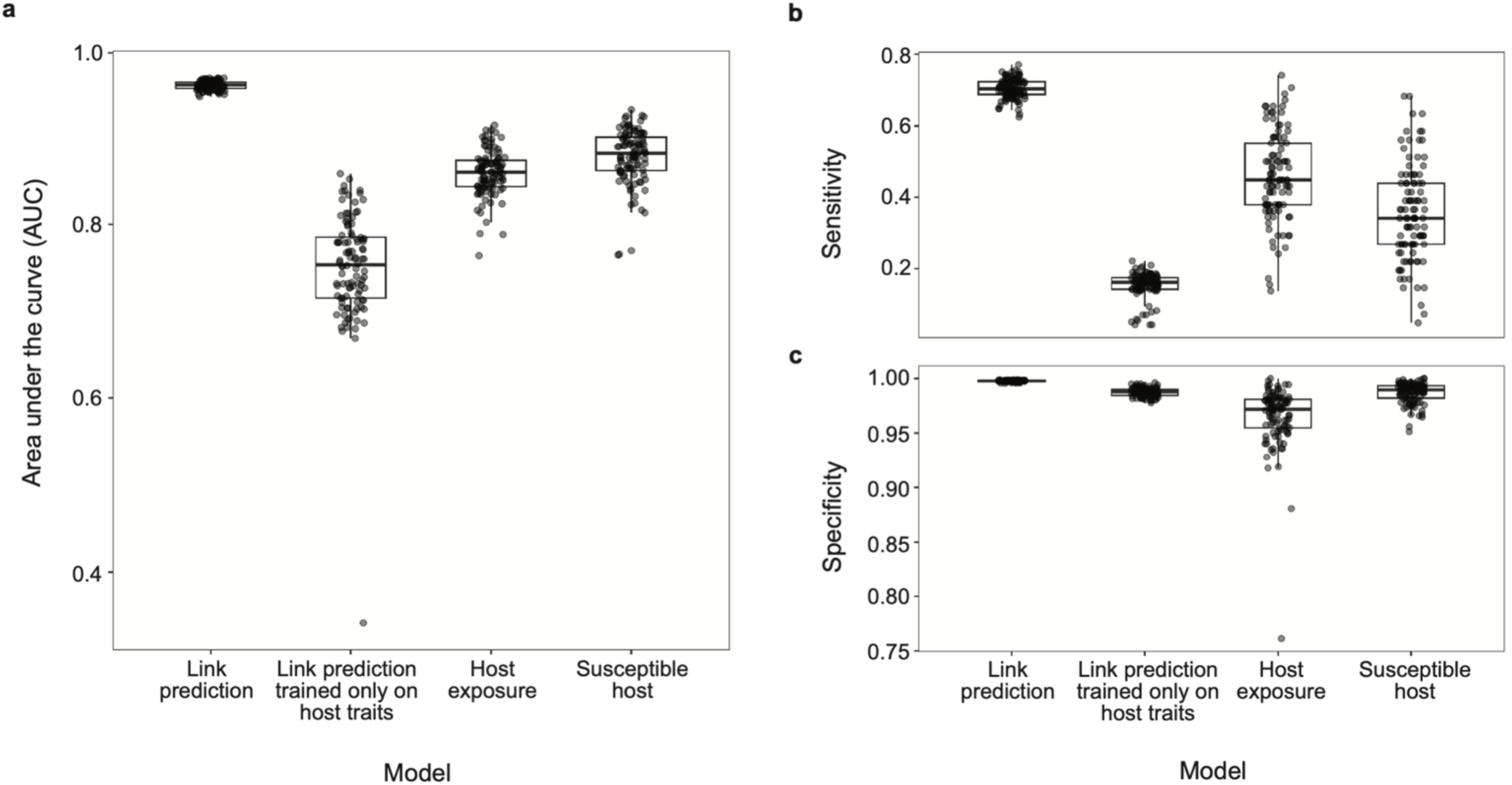
Performance measures of boosted regression tree models. Model accuracy was evaluated by (a) area under the receiver operating characteristic curve, (b) sensitivity and (c) specificity. Link prediction models were trained on known host-virus links as the response variable but differed in the inclusion versus exclusion of viral genomic traits as predictors. Host prediction models were trained on RT-PCR positivity (i.e., host exposure model) versus virus isolation data (i.e., susceptible host model). All models were trained on 100 random splits of training (70%) and test (30%) data.

**Fig 2.**
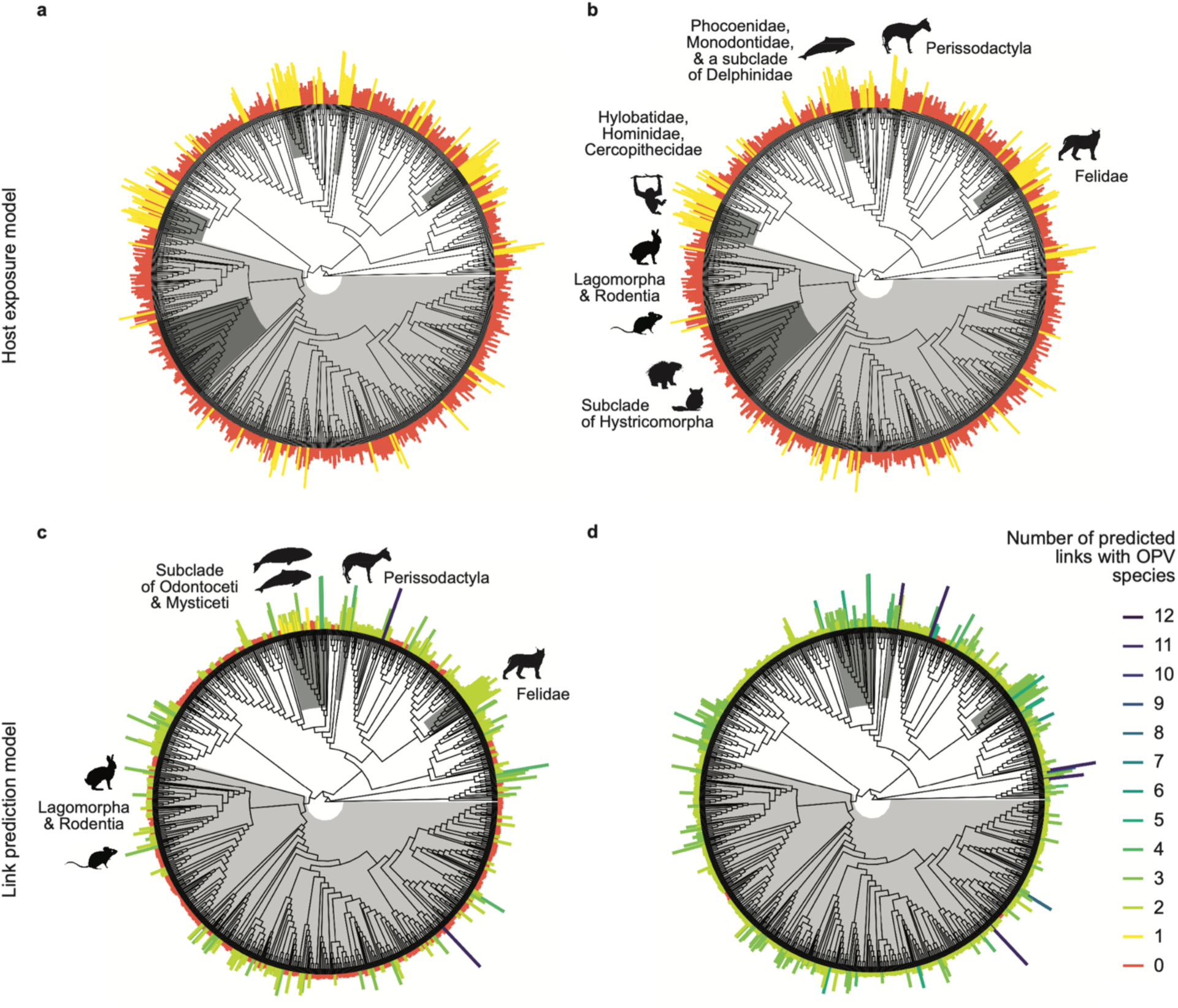
Predictions of *Orthopoxvirus* positivity reveal taxonomic patterns and the effects of threshold moving. For (a, b) the host exposure model and (c, d) the link prediction model, bars**/**segments are scaled by predicted probabilities and colored by binary classification of predicted host genera assuming an 80% sensitivity threshold (a, c) or a threshold maximizing the sum of sensitivity and specificity (b, d). For the link prediction model, if multiple links were predicted per host genus, their predicted probabilities were averaged to represent the genus’ mean probability of hosting any OPV. Clades identified through phylogenetic factorization with significantly different mean predictions are shaded in grey with corresponding labels and representative mammal silhouettes, which apply across panels of the same model: (a, b) for host exposure and (c, d) for link prediction

### Link prediction model

In contrast, the link prediction model trained on host traits and viral features (to predict the existence of a host-virus link) classified compatible host-virus pairs with higher accuracy, higher specificity, and higher, moderate sensitivity (Fig 1; S1 Table). To explore the robustness of the link prediction model, we tried predicting links solely on host traits, which, similar to host prediction models, resulted in lower model accuracy, specificity and sensitivity compared to training on both host traits and viral features (Fig 1; S1 Table). We also tried excluding host associations with vaccinia virus from our link prediction model trained on host and virus traits. This decision was based on the fact that multiple passages *in vitro* of vaccinia virus have led to the artificial selection of viral genes implicated in virulence, replication capacity, and host range, which may not reflect natural host-virus interactions. Excluding links with vaccinia virus significantly increased sensitivity (*t* = -10.06, *p* < 0.001) and lowered specificity (*t* = 2.99, *p* < 0.001) and overall model accuracy, albeit marginally (AUC = 0.95 vs. 0.96; *t* = 10.1, *p* < 0.001) (Fig 1; S1 Table).

Link predicted probabilities, represented as the mean probability of hosting any OPV per host genera, displayed strong phylogenetic signal (Pagel’s λ = 0.87). Similar to the host prediction models, phylogenetic factorization predicted Felidae (*n =* 17 tips) more likely to host OPV and Lagomorpha and Rodentia (*n =* 514) less likely to host OPV (Fig 2; S2-S4 Tables). The order Perissodactyla (*n* = 8) was also predicted more likely to host OPVs in agreement with the host exposure model (S2-4 Tables), while overlapping taxa of the order Cetacea and Ziphiidae were predicted more likely to host OPVs by both the link prediction and susceptible host models (Figs 2 and S1; S3 and S4 Tables).

### Optimizing classification

To address the problem of class imbalance (i.e., the unbalanced representation of hosts and non-hosts or links and non-links in our training datasets), we explored various optimal thresholding methods for generating binary host classifications from the predictions of our host and link prediction models. Unsurprisingly, the number of predicted mammalian hosts for OPVs was highly sensitive to the classification threshold, particularly for the link prediction model (Fig 2; S5 Table). Applying an 80% sensitivity threshold (*t_ReqSens80_ =* 0.36) versus a 90% sensitivity threshold (*t_ReqSens90_ =* 0.28) to host exposure model predictions resulted in a 2.3-fold increase in the number of predicted host genera (from *n* = 116 to 263) (S5 Table). The same thresholds selection method applied to link predictions (*t_ReqSens80_* = 0.13 vs. *t_ReqSens90_* = 0.05) resulted in a 3.6-fold increase in the number of predicted host genera (from *n* = 502 using 80% sensitivity to n= 1,791 when using 90% sensitivity), which had a similar effect to applying the threshold where sensitivity equals specificity (*t_Sens=Spec_* = 0.04; *n* = 1,864) (S5 Table). As evident across all models, though, lowering the threshold value maximized sensitivity, ensuring that fewer positive cases were missed, albeit at the expense of a higher false positive rate.

Threshold selection also led to changes in the taxonomic composition of previously unobserved, predicted mammal genera when grouped by mammal order. For the link prediction model, increasing the required sensitivity from 80% to 90%, which lowered the threshold value, led to an increased representation of rodents (from 34% to 42%) and a decreased representation of carnivores (from 23% to 17%), while the percentage of unobserved predicted genera from other mammal orders remained approximately the same: e.g., Cetartiodactyla (15-16%), primates (10-11%), and Eulipotyphla (7-9%).

### Feature importance

We observed some overlap in the mammal traits identified as predictive of OPV positivity and compatible host-virus pairs. Across host trait models and both link prediction models (trained on host and viral traits vs. host traits only), PubMed citations of host genera, as an indicator of sampling effort, and dispersal potential (i.e., the distance an animal can travel between its place of birth and its place of reproduction) were consistently important predictors (Fig 3; S2 and S3 Figs; S6 and S7 Tables). To determine whether host citation counts, as a proxy for sampling effort, confounds the relationship between host trait profiles and OPV positivity, we conducted a secondary set of BRTs that modeled citation counts as a Poisson response, and found that host traits did not predict study effort based on its classification performance (mean AUC = 0.5, which is not different from chance). The two host traits with the highest relative importance in the in the model including both the viral and host traits (Fig 3C), which had the best classification performance, were *island dwelling* (the proportion of species in a genus having 20% or more of the breeding range occurs on an island) and *dispersal* (median distance in km travelled by species in a genus, between the birth site and the breeding site). Both variables followed a positive association with the probability of hosting OPVs, although their effect sizes were small (S4 Fig).

**Fig 3.**
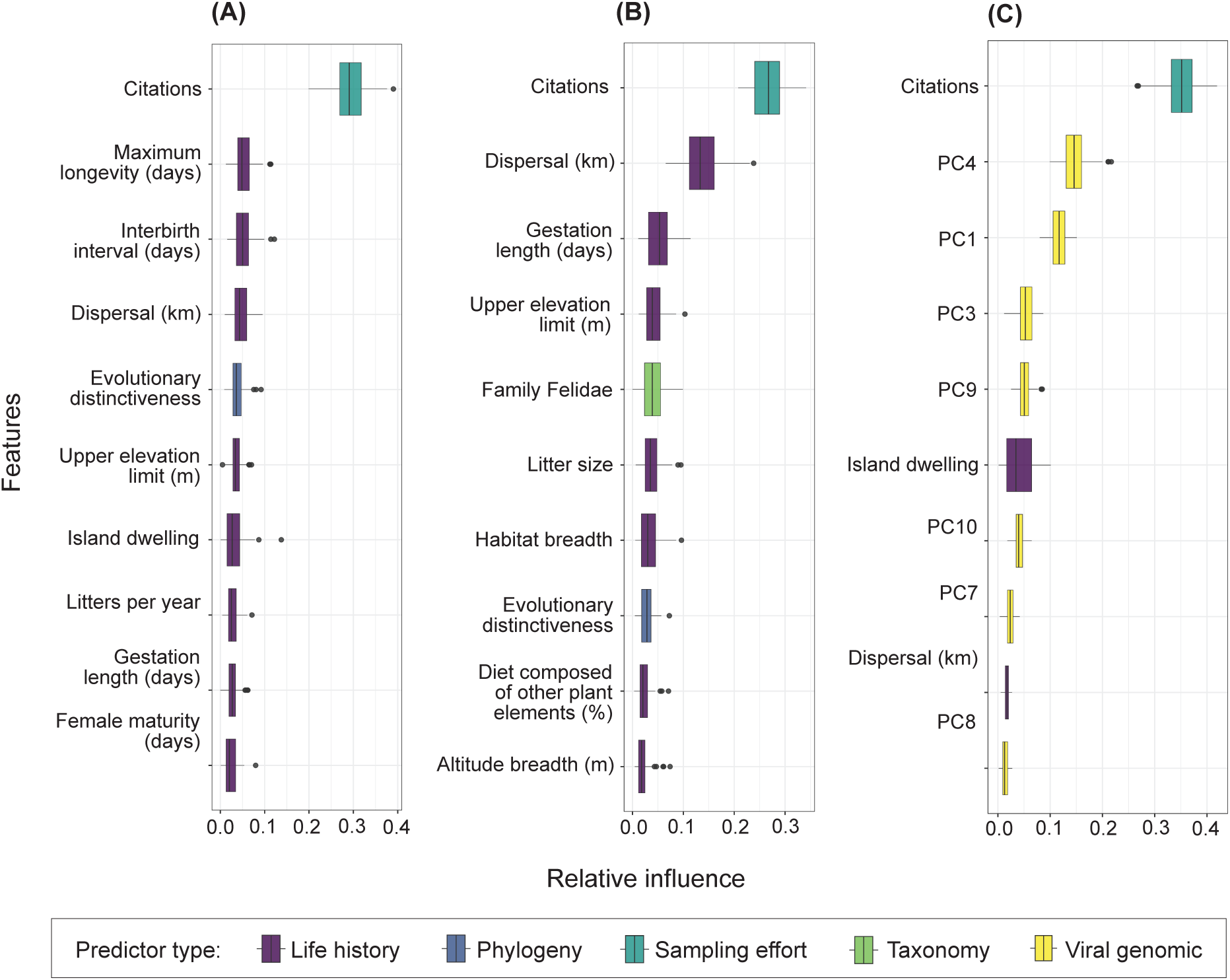
Relative influence of model features ranked. The most important predictor variables in BRT analysis are shown for (a) the host exposure model, (b) the susceptible host model, and (c) the link prediction model trained on a combination of host-virus traits, including the principal components (PCs) of viral accessory gene data, with explicit pairing of host-virus links as the response. Each horizontal bar plots the variability in the relative influence of the predictor variables as measured across 100 random partitions of training (70%) and test (30%) data. Boxplots show the median and interquartile range, whiskers are the extremes, and circles are additional outliers.

In our link prediction model trained on host and viral traits, viral features were represented by ten principal components (PCs) derived from a principal component analysis (PCA; Supporting Methods). These components captured the majority (70%) of the variance in the presence/absence of viral accessory genes (S5 Fig). While both host prediction models and the link prediction model trained solely on host traits ranked life history traits highly (Figs 3A and 3B; S6 Fig), rankings of relative feature importance for the link prediction model identified a multitude of PC variables as important predictors (Fig 3C). Further analysis of the influential PCs and their loadings, which indicate the relative contribution of individual genes to each PC, found potential patterns in the functional roles of genes with high positive or negative loadings (S1 Data). For instance, genes with the greatest contributions to PC4 and PC3 largely encoded proteins involved in host interaction and immune evasion, including non-essential membrane proteins and proteins that target innate cytokines/chemokine and cell death inhibitors (Fig 4; S7 Fig). Notably, for PC1, genes with the greatest positive loadings were predominately associated with cowpox virus, while genes with the greatest negative loadings were primarily associated with mpox virus (S1 Data), indicating clustering among specific OPV species (S8 Fig) that may not have been fully captured by the accessory genes included in the PCA. A list of NCBI accession numbers and PCA loadings can be found in the github repository (github.com/viralemergence/PoxHost/data).

**Fig 4.**
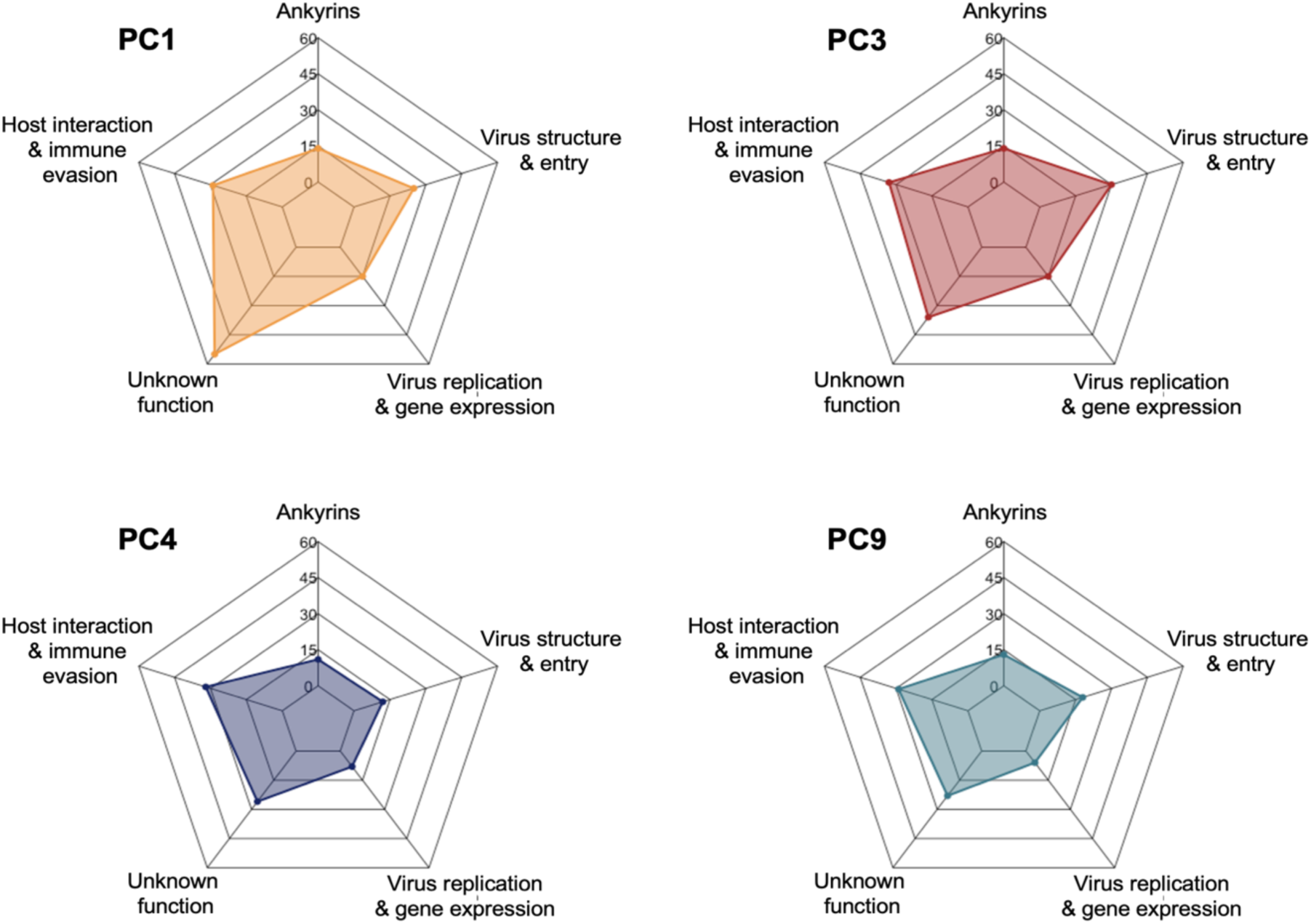
Relative contribution of predicted functional groupings among genes influential to principal components (PC) 1, 3, 4, and 9. For the most influential viral traits based on link prediction, we assigned predictive functions to genes with loading values greater than 1.5 times the standard deviation in the positive range or less than 1.5 times the standard deviation in the negative range (*n* = 89,128, 123, and 89, respectively). We then plotted counts for each gene category (ankyrins, virus structure and entry, virus replication and gene expression, host interactions and immune evasion or unknown function) for PC1, PC3, PC4, and PC9.

### Distribution of potential *Orthopoxvirus* hosts

Mapping of the geographic distribution of observed hosts (i.e., known links) alongside predicted host genera (observed and unobserved) revealed novel hotspots of overlapping OPV hosts in parts of Indonesia and Malaysia, southern East Africa, the West African coastline, the Amazon basin, and the Brazilian coastline (Figs 5A and 5B). We identified the highest concentration of potential OPV hosts in the Eastern Himalayas, western Cameroon, and regions of Central and Eastern Africa extending from the Democratic Republic of Congo to Kenya (Fig 5B). Similar comparisons of observed vs. predicted host genera for mpox virus revealed novel hotspots in the rainforested regions of Central and Eastern Africa and parts of the southern and northern US plains (S9A and S9B Fig). Our models show high overlap between geographic regions with the most potential OPV host species (Fig 5B) and areas where the smallest percentages of the population have been vaccinated against smallpox [26](Fig 5C) and are, therefore, at higher risk of zoonotic OPV emergence.

**Fig 5.**
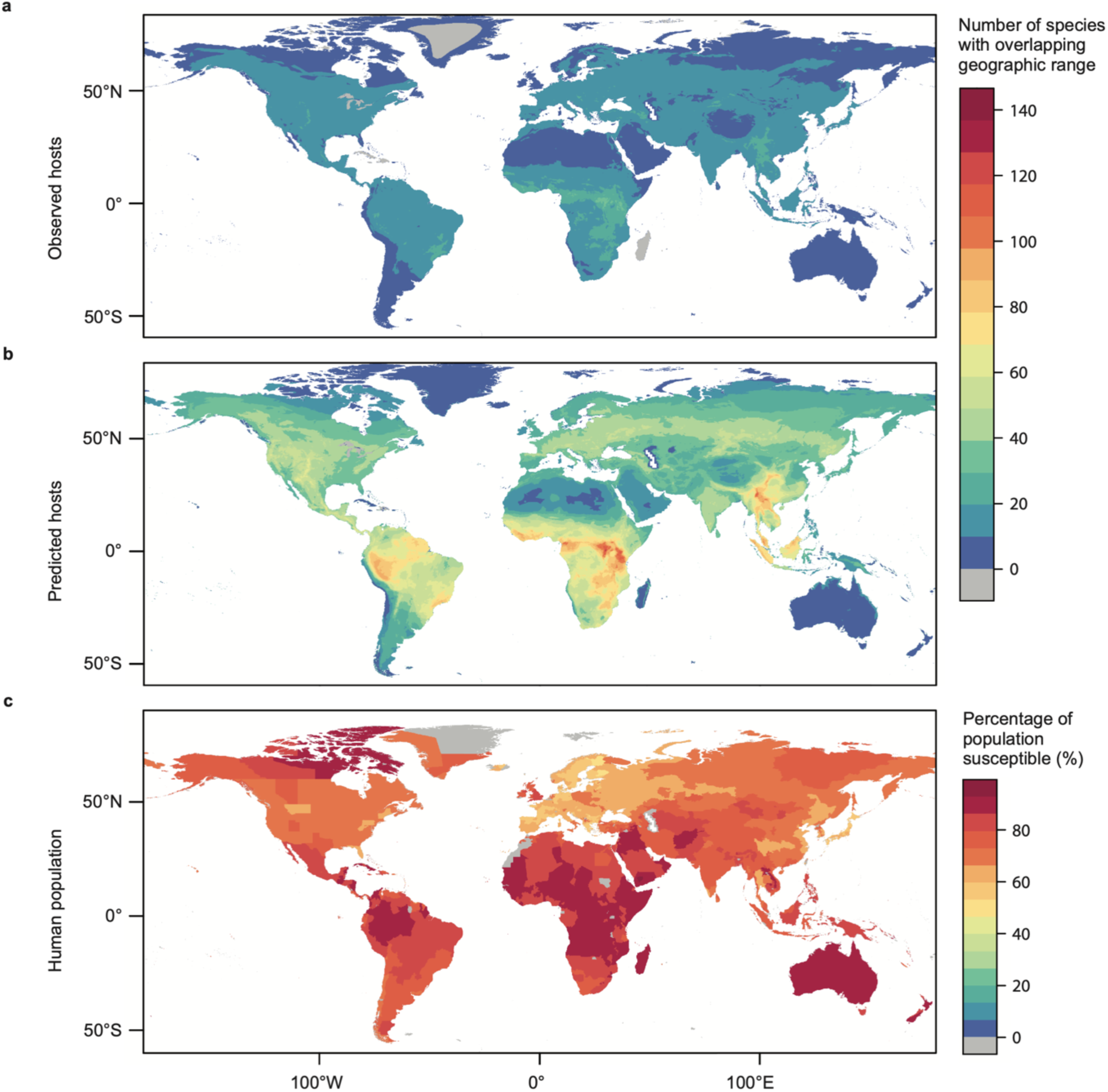
Geographic distribution of *Orthopoxvirus* hosts. Host distributions are based on the IUCN Red List database of mammal geographic ranges for those species belonging to (A) observed host genera and (a) predicted (observed and unobserved) host genera based on the results of the link prediction model and applying a 90% sensitivity threshold for host classification. The corresponding legend depicts the number of species with overlapping geographic range by color. (c) depicts smallpox vaccination coverage among humans (adapted from Taube *et al.* 2022), whereby the legend indicates the percentage of the population susceptible to orthopoxviruses.

## Discussion

Machine learning models that leverage ecology to predict likely reservoirs of emerging pathogens can be remarkably accurate and can be used to guide the selection of target host species for monitoring and surveillance. However, trait selection based on host ecology alone can overlook the potential influence of genetic diversity on the ability of pathogens to infect a broad or narrow range of hosts. Here, we focused our study on OPVs, a group of closely related viruses whose genetic variability is thought to contribute to their diverse host range. Using BRTs trained on observed associations between OPVs and mammal hosts, we found that a model informed by both viral genomic features and host ecological traits predicted mammal hosts with improved accuracy compared to a model trained on host traits alone. Moreover, variables capturing viral accessory gene data were found to be highly predictive of host-virus compatibility, illustrating how integrating viral genomic traits, such as the presence or absence of virulence genes, can improve host prediction and help identify key features associated with host specificity. Relating the most influential PCs in model prediction to their original features, we classified genes with the highest loadings by their predicted functional roles, which offers new opportunities to explore the roles of accessory genes in the prediction of OPV host range.

Genomic features such as phylogenetic relatedness and codon usage bias can be informative of host prediction [14,27,28] but are rarely used alongside host ecological traits to inform model prediction. Previous work has demonstrated the utility of ensemble modeling for a multi-perspective, host-virus-network framework in predicting viral host ranges by integrating multiple similarity learners derived from three perspectives: genomic (i.e., nucleotide bias, codon usage, secondary structure features, and genomic dissimilarity), ecological (mammalian traits), and network-based [29]. The network perspective utilized a bipartite graph of known virus-host interactions to quantify node similarities, enabling predictions for unobserved virus-host associations based on how viruses and hosts are positioned in the network. All three perspectives were combined via ensemble modeling to enhance predicting accuracy, but were originally modeled separately. Building on this foundation, we integrate genomic, ecological, and network-based perspectives into a unified model to address the underestimation of host-virus associations. This approach demonstrates how incorporating viral genomic features in link prediction can also serve as a valuable tool for public health surveillance and generate new hypotheses about host-virus compatibility. Our study found that BRTs trained on both host and virus traits produced similar and on average higher AUC, specificity, and sensitivity than BRTs trained solely on host traits alone. The architecture of our model was qualitatively similar to that of Blagrove et al. [17], which used a link prediction approach incorporating both host and viral traits to predict potential hosts of poxviruses. Despite differences in feature selection and methodology, their model, like ours, had low to moderate sensitivity, which was attributed to an imbalance in the distribution of observed versus unobserved host-virus associations. While Blagrove et al. corrected for class imbalance using over-sampling methods, we opted to explore the effects of probability threshold selection on host predictions, offering an alternative and more informative approach to addressing this challenge.

We note that in both our host and link prediction models, citation counts of host genera was highly ranked (Fig 3), reflecting the importance of sampling effort in accurate prediction, as genera with higher citation counts are more likely to have been studied for viral associations and have greater representation among OPV sequence data. A dataset rich in variation in both viral genomic features and host species improves generalizability and out-of-sample prediction of the model. We note that the family Felidae is well-represented as hosts across diverse cowpox virus sequences which may contribute to the high predicted probability of detecting OPVs in the family Felidae. Nearly half of all represented OPV genomes originated in humans. In this dataset, mpox virus, cowpox virus, and vaccinia virus are relatively well-represented while borealpox virus and cetaceanpox virus are the least represented with a single host and viral genome. We included citation counts as an explanatory variable to take into consideration the impact of imbalanced sampling effort across taxa.

The problem of class imbalance is a common challenge when predicting the host range of emerging pathogens. Not only is there limited data on rare or novel hosts, but also natural variation in species abundance and host susceptibility along with sampling bias can lead to imbalance in the representation of hosts and non-hosts (i.e., compatible and non-compatible host-virus pairs). When class imbalance is severe, applying a default threshold of 0.5 will often result in poor model performance. Thus, handling class imbalance is an important step in predictive modeling. In our study, we elected to explore higher sensitivity thresholds (e.g., 80% and 90% sensitivity) to reduce the risk of false negatives (i.e., potential hosts that are incorrectly classified as non-hosts). In the context of zoonotic host prediction, false negatives can lead to missed opportunities for surveillance and limit our understanding of the diversity of the zoonotic risk landscape for various OPV species. Thus, this trade-off ensures that fewer true hosts are overlooked, making our predictions more useful for proactive and precautionary measures in public health and zoonotic disease surveillance. Albeit a simple approach, tuning of the threshold parameter can have large implications on the interpretation of model predictions (Table S5). For instance, our PCR host exposure model trained on host traits alone estimated a 2.0 to 4.5-fold increase in the number of potential OPV hosts, assuming an 80% and 90% sensitivity threshold. Applying the same thresholds to our link prediction model trained on host and viral traits resulted in a 5.0 to 17.7-fold increase in the number of potential hosts, a likely artefact of the large proportion of pseudoabsences that construct our link prediction dataset. This severe class imbalance yielded lower predicted probabilities across BRTs making our ensemble of classifiers highly sensitive to threshold selection as compared to BRTs trained to predict OPV positivity on host traits alone.

Given the wide range of predictions that can result from threshold selection, choosing the optimal thresholding method should depend on the goal of the model and the context of the predictive task. For instance, borealpox virus is a recently emergent OPV only identified in humans thus far. While no formal surveillance data has been published, bulletins including communications from the CDC have implicated voles as a putative reservoir. For the purpose of this study, we proceeded with only the human host data. Based on a restrictive 80% sensitivity threshold, our link prediction model predicted two potentially previously unobserved host genera, *Ailurus* (red panda) and *Micromys* (harvest mice), neither of which have a geographic range overlapping with known zoonotic events, and are thus, unlikely current reservoirs in the observed virus geographic distribution. However, applying a less restrictive threshold of 90% sensitivity resulted in the prediction of eight potential previously unobserved hosts for borealpox virus, including the only known reservoirs of ectromelia virus, rodents in the cosmopolitan genus *Mus* (mice). Notably, genomic analysis of borealpox virus suggests a history of recombination with ectromelia virus, including in the putative gene encoding the A-type inclusion protein [30]. Deletion of the A-type inclusion protein in cowpox virus enhances viral replication in *Mus musculus* [31], suggesting that variants or deletion mutations in this gene contribute to species specificity. We suspect this recombination site drove the indicator for *Mus* species as a potential borealpox virus reservoir in our model. In recapitulating the ectromelia virus recombination site with borealpox virus, our findings suggest that when sample data are sparse, applying less restrictive thresholds may be necessary to reveal informative, meaningful predictions by identifying a broader set of potential hosts for guiding targeted surveillance of a rare or novel virus. Conversely, when sample data are abundant, a more restrictive threshold may be appropriate to identify a manageable number of potential hosts and prioritize specific candidates for detailed investigation.

Similarly, threshold selection can also impact the taxonomic representation or composition of model predictions. For instance, among known mammal genera susceptible to mpox virus, the majority are from the Rodentia (37%) and Primate (37%) orders, which is consistent with Blagrove et al. whose findings predicted a similar composition of mpox virus hosts (80% from the Rodentia and Primate orders) [17]. However, we show in our link prediction model, that the host composition of model predictions broadens when applying a less restrictive threshold. When assuming an 80% sensitivity threshold, 45.2% of previously unobserved mpox virus hosts were predicted to be from Rodentia, while increasing model sensitivity to 90% led to a decrease in predicted rodent genera (31.4%) and a substantial increase in genera from the Carnivora order (from 3.2% to 19.6 %), particularly those in the Felidae (cats), Canidae (canids), Mephitidae (skunks), Mustelidae (mustelids), and Procyonidae (raccoons) families. Thus, choosing the optimal threshold for classification may have important public health implications, as it may expand the focus on species that may be overlooked when applying a more restrictive threshold. In the mpox example, this is of particular interest since its recent global spread outside endemic countries raises concerns of the potential for transmission from humans to wildlife or spillback in nonendemic areas [32] and the establishment of new animal reservoirs. We note that the genus *Rattus* was not predicted as a likely host for mpox virus in our link prediction model at any of our tested thresholds which is consistent with laboratory experimental data showing that rats are not susceptible to mpox virus and in contrast to predictions using different approaches [17]. A table of predicted binary host-virus links can be found in the github repository (github.com/viralemergence/PoxHost/figures/other/linkpred).

Our study reveals a striking correspondence between regions with high potential OPV host species diversity and those where smallpox vaccination rates are lowest (Fig 5). This juxtaposition suggests a pronounced vulnerability to zoonotic OPVs within these populations. Hotspots identified as having high concentrations of postulated OPV hosts included several distinct areas within the eastern Himalayas of South Asia bordering southwest China, Central and Eastern Africa, Indonesia, and Malaysia – areas where wildlife sampling and monitoring of spillover events should focus their efforts. Comparing the geographic distribution of predicted hosts to known hosts also revealed potentially under sampled areas along the West African and Brazilian coastline; these areas also coincide with many recent human cases of reemerging zoonotic OPVs including mpox virus in Cameroon [33] and vaccinia virus in Brazil [34]. Interestingly, greater geographic dispersal (distance traveled between birth site and breeding site) was not only an important predictor (Fig 3) but was also positively associated with an increased probability of hosting OPVs (S2, S3 and S4 Figs), supporting theories of host dispersal as a key contributor to the evolution of host-pathogen dynamics [35]. Specifically, as individuals move to new areas to breed, they must adapt to new environmental conditions and, in the process, are exposed to and may host more diverse pathogens. On the other hand, the positive association between island dwelling and the probability of hosting OPVs could be indicating higher prevalence or oversampling in constrained island communities, although the effect size was small (S4 Fig).

Like all related studies, our study had several limitations, particularly with respect to sampling bias, incomplete datasets, and model interpretability. First, in zoonotic host prediction, host-virus association data is inherently incomplete, a limitation further compounded by sampling bias, as host-virus interactions are disproportionately observed in well-studied taxa and regions, especially for human-associated OPVs. Although we incorporated pseudoabsences to enhance generalizability, our predictions may overemphasize well-documented interactions, while underestimating associations involving rare or poorly sampled hosts, or novel human infecting viruses. Ongoing efforts to consolidate host-virus interaction data will help to reduce biases and improve data quality over time.

Second, as only complete OPV genomes were extracted from NCBI for inclusion in the link prediction model, the number of host-virus associations with accessory gene data available was limited. Moreover, missing data for host and virus traits precluded more precise predictions at the species level. Thus, we aggregated the dataset at the genera level to reduce the number of pseudoabsences in the models. We expect future prediction models to improve in their accuracy as whole genome sequences of host and virus become more readily available. Third, our study elected to classify accessory gene data based on presence-absence, trading simplicity over potential loss of information that could have related protein activity to host-virus compatibility. Subsequent studies could investigate transforming accessory gene data as continuous variables based on amino acid conservation or ranked categorical variables, where accessory genes are missing, truncated and non-functional, truncated but likely functional, or intact, as previous studies have explored [36]. Reducing accessory gene variables to their principal components (to avoid overfitting our model) also precluded us from interpreting the effects of individual accessory genes in predicting host-OPV pairs. However, our findings could inform future iterations by training a model on the genes identified in this study as contributing highly to influential principal components, thereby directly exploring the relationship between specific viral accessory genes and host range. Furthermore, many functional roles of genes were predicted based on sequence homology to known OPV accessory genes, particularly for vaccinia virus or mpox virus, but are not well characterized in other species. Because there is no standard for naming genes, many functions are putative and gene names can be based off other viruses, which can be misleading. Finally, predictive models can also be limited by the machine learning algorithms they employ. As black box models, BRTs can be difficult to interpret and do not provide a complete or exact understanding of the relationships and potential interactions between predictors and the response variable. While our study employed cross-validation to assess the model’s robustness, we did not perform external validation or biological validation, which requires field and laboratory studies to confirm predicted host-virus interactions. Nevertheless, our model can also provide guidance for empirical sampling and initial insight into the molecular signatures of host-virus compatibility. While not the aim of our study, future link prediction models informed by viral genomic data could investigate training on different types of infection evidence to gain molecular insights into host capacity.

Historically, zoonotic OPVs like mpox virus were associated with small, sporadic outbreaks limited to endemic countries. However, large outbreaks in recent years including the worldwide spread of mpox virus in 2022 warrant growing concerns about the potential for mpox virus and other OPVs to establish new endemic areas. Recent evidence of animal-to-human and human-to-animal transmission of mpox virus also suggest the potential for new reservoir species to exist in traditionally non-endemic regions, underscoring the importance of predicting the host range of emerging pathogens. Adapting models to predict compatibility is also crucial to expanding the growing toolkit of analytical models used to predict host-pathogen interactions. Here, we demonstrate how trait-based machine learning models can be trained on both mammal traits of potential hosts and the genomic features of OPVs to improve the accuracy of host prediction and gain a more comprehensive understanding of the factors that influence host-virus compatibility. By exploring the trade-off between sensitivity and specificity through classification threshold selection, we show the importance of aligning models to meet the specific needs and objectives of host prediction for greater practical application.

## Materials and methods

We used OPVs as a case study to develop and validate a multi-perspective modeling framework for the prediction of host-virus associations. We generated two types of trait-based models to compare model performance and predictions: (1) a host prediction model trained separately on two evidence levels for host capacity (i.e., host exposure via PCR positivity data and host susceptibility via virus isolation data) using host ecological traits alone; and (2) a link prediction model trained on explicit pairing of host-virus associations using a combination of host and virus features. The former model type predicted host positivity for OPVs, a group of viruses, while the latter predicted the existence of a host-virus link. Our modeling targets were to identify taxonomic and phylogenetic patterns in the predicted reservoir hosts of OPVs along with their corresponding geographic distribution, rank influential host and viral features, and investigate the functional roles of accessory genes that contributed substantially to link prediction.

### Datasets

For the host prediction model, we obtained known host positivity data from the Global Virome in One Network (VIRION), an open database of host-virus interactions drawn from scientific literature and online databases [37]. We included only host positivity detected via PCR (molecular detection) or virus isolation and excluded detections with variola virus, as the virus was eradicated prior to the development of many modern diagnostics. We then collapsed host positivity data to the genus level and merged the data with the broader mammal taxonomy to obtain pseudoabsences for mammal genera with unobserved OPV associations [38]. Next, we derived host predictors from COMBINE, a published mammal dataset of 54 morphological, biogeographic, and life history traits that is taxonomically compatible with the IUCN Red List v. 2020-2 and PHYLACINE v. 1.2 [39–41]. We applied summary measures to represent ecological trait data at the genus level by transforming all categorical variables into multiple binary variables. We then aggregated binary variables to the genus level assuming the mean to obtain the proportion of species in a genus having the variable outcome/trait, and we aggregated continuous and integer trait variables assuming the median. To account for large gaps in trait coverage, we excluded variables with zero variance or with data for less than 60% of host genera (S10 Fig), resulting in 61 ecological trait variables. We also generated binary predictors for each mammal family to represent taxonomy, leading to an additional 42 taxonomic variables in the host trait model. Lastly, we incorporated a variable for evolutionary distinctiveness (*ed_equal*) using the *picante* package in R and a count variable of the number of scientific publications on each host genera (*cites*) as a measure of sampling effort using the R package *easyPubMed.* A complete list of host predictors incorporated in the host prediction model can be found in the supporting information (S6 Table).

For the link prediction model, we merged data from VIRION with host-OPV associations obtained from NCBI by extracting OPV genomes and their annotations (https://www.ncbi.nlm.nih.gov/)(S2 Data). We included viral isolates with full genome sequences from experimental laboratory studies when possible. Due to the abundance of available mpox virus and cowpox virus genomes, we included a subset of each for all genomes with novel host associations or representing divergent clades. Multiple sequences for a particular virus-host combination were only included if an annotated genome was available and there were differences in the presence/absence of accessory genes. Again, we merged host-virus association data with the broader mammal taxonomy to accommodate out-of-sample predictions, expanding our mammal-virus network to include all combinations of mammal genera and OPV species. As predictors in our model, we incorporated the same host predictors as in the host prediction model, with the exception of 110 taxonomic variables instead of 41. Additionally, we incorporated viral predictors by identifying accessory genes from the aforementioned OPV genomes using Roary (https://sanger-pathogens.github.io/Roary/) with 80% minimum sequence identify for blastp and 95% of isolates a gene must be in to be designated as part of the core genome. After transforming the accessory gene data into a binary matrix across 981 accessory genes from 197 OPV sequences, we conducted PCA to reduce the dimensionality of our dataset and distill presence/absence variables down to their most important features. The first ten principal components, which explained roughly 70% of data variance, were then included as viral predictors in our link prediction model (S5 Fig). OPV genomic data used in the PCA was available for 12 OPV species, but only for a limited number of OPV-host pairings (e.g., Mpox virus-human, Mpox virus-dog). However, as we considered the 10 PC variables derived from the PCA as viral traits, we conducted median imputation of the PC values for the links between hosts with any of those 12 OPV species for which PC data were missing (e.g., for all unobserved Mpox virus-host pairings we applied the median PC value for Mpox virus, for each one of the PCs). A complete list of host and viral features incorporated in the link prediction model can be found in the supporting information (S7 Table).

Lastly, for phylogenetic analysis, we used a supertree of extant mammal species trimmed to the genus level [38] (https://vertlife.org/data/mammals/). Data collation (cleaning and merging) were conducted in R version 4.2.2. Additional details regarding the cleaning of final datasets used in the host trait and link prediction models are available in the supporting information including dimension reduction for generating viral predictors and taxonomic reconciliation.

### Statistical analysis

#### Host prediction models

Following Mull et al. [42] (https://github.com/viralemergence/hantaro), we used a trait-based model to infer the host range of OPVs, integrating only host mammal traits as predictors. This approach handles binomial virus positivity of host genera as the response variable and host trait data as predictors. It uses a machine learning algorithm to identify host features associated with OPV detection and to make predictions of the probability of OPV detection based on how similar a species’ trait profile is to that of observed host species. False positive results, where a species classified as a pseudoabsence is predicted to host a virus, are used to infer potential, previously undetected or unobserved host genera.

To classify mammal genera as OPV hosts based on our matrix of predictors/traits, we used boosted regression trees (BRTs). BRTs combine two validated machine learning algorithms: regression trees, which model the relationship between outcome and predictors by recursive binary splits, and boosting, an adaptive technique in which the model considers the prediction errors of previous trees when fitting subsequent trees thereby improving predictive performance and classification accuracy [43]. BRTs are advantageous to our study as they can handle large ecological datasets, a binomial outcome variable, and a diversity of model predictors and the potential interactions between them. They can also fit potentially complex nonlinear relationships, produce response functions for each predictor, and are robust to missing values and outliers [44].

Using BRTs, we developed separate evidence type models for classification of virus positivity: a host exposure model based on PCR-detection and a susceptible host model based on virus isolation data. Prior to model fitting, we used the R package *rsample* to randomly partition our data into training (70%) and test (30%) sets and apply stratified sampling to ensure an equal distribution of positive labels in both datasets. Next, we tuned our model by training it on multiple combinations of hyperparameter values allowing for a maximum number of 5,000 trees, an interaction depth ranging from 2 to 4 (to control tree complexity), a shrinkage (learning) rate ranging from 0.0005 to 0.01, and a minimum number of 4 observations in terminal nodes (S11 Fig). Using the *gbm* package in R (Greenwell et al., 2020) for all model fitting, we applied five-fold cross validation and a bag/subsampling fraction of 0.5, and we assumed a Bernoulli distribution error for our binary response variable. Based on multiple model performance measures, we selected the best set of parameter values for model training, which included a learning rate of 0.01, and an interaction depth of three and the maximum number of trees set to 4500. With our tuned model for prediction, we set citation counts per genera to the mean across all mammal genera to adjust for sampling effort. We then fit replicate models to 100 randomly stratified partitions creating an ensemble of BRTs for each evidence type (PCR and virus isolation). Additionally, we constructed a third set of BRTs to determine the influence of sampling effort on the trait profiles of OPV positive genera; instead of virus positivity, we modeled the citation counts of host genera as a Poisson response variable applying the same optimal hyperparameter values except for 10,000 maximum trees.

#### Link prediction model

To predict the host ranges of multiple OPV species and to allow for integration of viral genomic features, we adapted the host prediction models l for link prediction whereby the model predicts the existence of a host-virus link (as opposed to positivity for a virus or group of viruses). While the previous host prediction models predicted OPV positivity (regardless of OPV species), the link prediction model uses explicit pairing or ‘links’ between host genera and virus species as a response variable and, therefore, allows for the inclusion of both viral and host traits simultaneously as predictors. As in the host prediction models, undiscovered host-virus associations are identified based on false positive predictions.

Using BRTs to predict the probability of links (associations between host genera and virus species), we followed a similar workflow as the host prediction models for generating link predictions. We checked that both test and training partitions contained similar proportions amongst the included viral taxa. We developed three ensembles of BRTs. The first, which included all combinations of host genera and OPV species (as described in *Host-virus association data* of S1 Text), used a combination of host and viral features to predict host-virus links. The second used only host features for prediction. The third used both host and viral features for prediction but excluded any host-virus combination with vaccinia virus. The latter was intended to explore if spontaneous, extended deletions and other modifications in viral genes implicated in virulence, replication capacity and home range of vaccinia virus due to multiple passages in cell cultures, may introduce bias in the genetic features included in the model and affect model performance.

#### Model performance and prediction

For each model fitting of the partitioned training and test data, we obtained model performance metrics including AUC, sensitivity and specificity using the *ROCR* and *InformationValue* packages in R and a threshold value of 0.5. For each ensemble model, we then calculated the mean and standard error of each performance metric measured across the 100 random partitions and conducted an unpaired *t*-test to compare model performance, with *p*-values adjusted for the false discovery rate using the Benjamini-Hochberg correction [45,46].

To compare model predictions, we first calculated the Spearman correlation coefficient between the predicted probabilities of the host exposure vs. susceptible host models as well as the link prediction model when including vs. excluding vaccinia virus. Next, we assessed model predictions for taxonomic patterns. Because the link prediction model predicted multiple links per host genera, we calculated mean predictions per host genera to obtain a single value per tree tip (branch endpoint) for phylogenetic analysis. Using the *nlme* package in R, we estimated host phylogenetic signal based on Pagel’s λ, a measure of the degree to which the propensity for virus positivity among related genera is driven by shared evolutionary history. We then identified clades with significantly different propensity for hosting OPV at various taxonomic resolutions using phylogenetic factorization, a graph-partitioning algorithm that explores the differences in measured traits between pairs of taxa while accounting for phylogenetic dependencies [47]. To determine the significant number of phylogenetic factors (clades), we adjusted for the family wise error rate using Holm’s sequentially rejective approach with a 5% threshold [48]. Phylogenetic factorization was implemented using the *phylofactor* package in R.

To generate binary predictions of OPV hosts and non-hosts, we transformed predicted probabilities into binary classifications using the *presenceabsence* R package. Based on the previously described ROC, we selected multiple thresholds to generate predictions based on different methods of threshold optimization. First, we explored higher sensitivity thresholds to prioritize minimizing false negatives (i.e., potential hosts that are incorrectly classified as non-hosts) at the expense of increasing the number of false positives (i.e., mammal genera incorrectly predicted to be hosts). We specifically selected an 80% sensitivity threshold (such that 80% of observed links are detected) and a 90% sensitivity threshold, as examples of more and less restrictive thresholds, or higher and lower threshold values, respectively. Second, we quantified the sensitivity of results to the choice of threshold by calculating the number of predicted hosts at additional threshold levels based on sensitivity-specificity trade-offs. These additional approaches included finding the threshold where sensitivity equaled specificity and findingthe threshold that maximized the sum of sensitivity and specificity, otherwise known as the Youden Index. We then compared classifications across models and demonstrated the effects of threshold selection by plotting predicted probabilities and their binary classifications on a circular phylogenetic tree using the package *treeio* and *ggtree*. Lastly, we mapped the geographic distribution of predicted observed and unobserved mammal genera; using the IUCN Red List database of known ranges of mammal species, we layered shape files of species belonging to thresholded genera to identify and contrast geographic patterns in model predictions.

Finally, we ranked model features by their mean variable importance (i.e., relative influence coefficients) and assessed similarities between models using the Spearman rank correlation coefficient, a rank-based measure of association. Viral PC predictors with high mean variable importance in our link prediction model were further analyzed by gene loading contribution to identify potential patterns in the functional roles of genes with high loadings. Specifically, we assigned predictive functions based on sequence homology to genes with loading values that were greater than 1.5 times the standard deviation in the positive range or less than 1.5 times the standard deviation in the negative range (S1 Data).

All statistical analyses were conducted in R version 4.2.2 and Microsoft Excel version 16.69.1.

## Author Contributions

K.K.T, P.F. and S.N.S. designed research; K.K.T., H.K., M.P.F. and S.N.S. performed research; D.J.B., R.G. and C.J.C. contributed to analytic tools; K.K.T, H.K., M.P.F. and S.N.S. analyzed data; and K.K.T., H.K., M.P.F. and S.N.S. wrote the paper. All authors reviewed the paper.

## Classification

Biological Sciences and Ecology.

## Data availability

Data used in this study is available as described in the methods section including the VIRION database (https://github.com/viralemergence/virion), COMBINE: the coalesced mammal database of intrinsic and extrinsic traits (Soria et al. 2021), global smallpox vaccination coverage (Taube et al. 2023). Accession numbers for OPV genomes included in this study can be found in Supplementary Table 2 and are accessible through GenBank (www.ncbi.nlm.nih.gov).

## Code availability

The R code to reproduce the analyses is available from the GitHub repository: https://github.com/viralemergence/PoxHost.

## Acknowledgments

This research was supported by funding to Verena (viralemergence.org) from the U.S. National Science Foundation, including NSF BII 2021909 and NSF BII 2213854. S.N.S. was partly supported by Washington State University and the US National Institute of Allergy and Infectious Disease/National Institutes of Health (NIAID/NIH) grant number U01AI151799 through the Centre for Research in Emerging Infectious Diseases—East and Central Africa (CREID-ECA). We thank three anonymous reviewers for their thoughtful critiques which have improved the quality of this manuscript.

## Supporting information

**S1 Fig. Predictions of *Orthopoxvirus* positivity for the susceptible host model trained on virus isolation data.** Bars/segments are scaled by probabilities and colored yellow according to binary classification of predicted host genera assuming (a) an 80% sensitivity threshold versus (b) a threshold maximizing the sum of sensitivity and specificity. Clades identified through phylogenetic factorization with significantly different mean predictions are shaded in grey.

**S2 Fig. Trait profiles of *Orthopoxvirus* positive mammal genera for the host prediction models**. Partial dependence plots for the ten most significant predictors across BRTs (applied to 100 random splits of training and test data) are displayed in order of relative importance for (a) the host exposure model trained on PCR data, and (b) the susceptible host model trained on virus isolation data as the response variable. Grey lines or points represent the marginal effect of each variable on predicting host status for each data split, while black lines indicate the average marginal effect. Histograms and rug plots illustrate the distribution of continuous and categorical predictors, respectively, across all included mammal genera.

**S3 Fig. Trait profiles of mammal genera with predicted existence of an *Orthopoxvirus* link for the link prediction model.** Partial dependence plots for the top ten predictors across BRTs (applied to 100 random partitions of training and test data) are displayed in order of relative importance for the link prediction model trained on (a) host and viral features and (b) host traits only. Grey lines or points represent the marginal effect of each variable on predicting host-virus links for each data split, while black lines indicate the average marginal effect. Histograms and rug plots display the distribution of continuous and categorical predictors, respectively, across all included mammal genera.

**S4 Fig. Marginal effects of the host traits with highest relative importance in the link prediction model when trained on host and viral features.** Partial dependence plots for island dwelling (A) and dispersal (B) across BRTs applied to 100 random partitions of training and test data are displayed in order of relative importance.

**S5 Fig. Cumulative variance plot for the first ten principal components (PC).** The dashed line displays the cutoff where the principal components capture approximately 70% of the variance in the viral accessory gene data.

**S6 Fig. Relative influence of model features ranked for the link prediction model trained only on host traits.** Each horizontal bar plots the variability in the relative influence of the predictor variables as measured across 100 random partitions of training (70%) and test (30%) data. Boxplots show the median and interquartile range, whiskers are the extremes, and circles are additional outliers.

**S7 Fig. Plot of *Orthopoxvirus* accessory gene loadings by the predicted gene function for each principal component (PC).** Loading plots with variables (i.e., genes) grouped by their predicted gene function are displayed for (a) PC1 and PC2, (b) PC3 and PC4, (c) PC5 and PC6, (d) PC7 and PC8, and (e) PC9 and PC10. Only genes with loading values greater than or less than 1.5 times the standard deviation from the mean are represented. Genes predicted to play a role in host interaction and immune evasion include those predicted to encode the Ankyrin family of proteins, Kelch-like proteins, and proteins involved in chemokine/cytokine regulation, cell death/cell cycle regulation, pathogen recognition, and antagonizing host adaptive immunity. **S8 Fig. Plot of *Orthopoxvirus* sequences by viral species for each principal component.** Score plots with samples (i.e., *Orthopoxvirus* genome) grouped by viral species are displayed for (a) PC1 and PC2, (b) PC3 and PC4, (c) PC5 and PC6, (d) PC7 and PC8, and (e) PC9 and PC10.

**S9 Fig. Geographic distribution of mpox virus hosts.** Host distributions are based on the IUCN Red List database of mammal geographic ranges for those species belonging to (a) observed host genera and (b) predicted (both observed and unobserved) host genera based on the results of the link prediction model and applying a 90% sensitivity threshold for host classification. The corresponding legend depicts the number of species with overlapping geographic range by color.

**S10 Fig. Host trait coverage across mammal genera.** Features with at least 60% coverage (denoted by the dashed line) across mammal genera were included in the BRT models. A complete list of feature coverage is available on the GitHub repository.

**S11 Fig. Performance measures of boosted regression tree models during parameter tuning.** Performance was evaluated by the area under the receiver operating characteristic curve (AUC), sensitivity, and specificity for (a) the host exposure model trained on RT-PCR data, (b) the susceptible host model trained on virus isolation data, and (c) the link prediction model during parameter tuning. Boxplots show the median and interquartile range alongside raw data for all 10 random splits of training (70%) and test (30%) data for each combination of learning rate and interaction depth.

**S1 Table. Mean performance measures for each ensemble model and their corresponding standard errors (SE).** Performance was evaluated by the area under the receiver operating characteristic curve (AUC), the sensitivity, and the specificity for each model assuming a threshold value of 0.5. We used 100 random partitions to generate an ensemble. An unpaired *t*-test with *p*-values adjusted for the false discovery rate using the Benjamini Hochberg correction compares the performance of the host exposure model (based on molecular detection of viral DNA using PCR techniques) to the susceptible host model (based on virus isolation from hosts) and that of the link prediction models with vs. without vaccinia virus associations. The effect size for the *t*-test was calculated using Cohen’s *d,* which standardizes the mean difference.

**S2 Table. Phylogenetic factorization of mean predicted probabilities for *Orthopoxvirus* positivity for the host exposure model.** The number of retained clades after a 5% family-wise error rate, taxa corresponding to those clades, number of species per clade, and mean predicted probabilities for the clade compared to the paraphyletic remainder are shown.

**S3 Table. Phylogenetic factorization of mean predicted probabilities for *Orthopoxvirus* positivity for the susceptible host model.** The number of retained clades after a 5% family-wise error rate, taxa corresponding to those clades, number of species per clade, and mean predicted probabilities for the clade compared to the paraphyletic remainder are shown.

**S4 Table. Phylogenetic factorization of mean predicted probabilities for host-*Orthopoxvirus* associations for the link prediction model.** The number of retained clades after a 5% family-wise error rate, taxa corresponding to those clades, number of species per clade, and mean predicted probabilities for the clade compared to the paraphyletic remainder are shown.

**S5 Table. Estimation of threshold values (*t*) based on different optimal thresholding methods.** Changes in the number of predicted hosts (*n*), sensitivity, and specificity were calculated for each threshold value applied to each ensemble model.

**S6 Table. Importance and ranks of mammal traits for the host prediction models.** The host exposure model (based on molecular detection of viral genomes using PCR techniques) and the susceptible host model (based on virus isolation from hosts) were trained on 105 predictor variables, including but not limited to 61 ecological features and 42 taxonomic trait variables.

**S7 Table. Importance and rank of mammal traits and viral features for the link prediction model.** The link prediction model was trained on 183 predictor variables, including but not limited to 10 viral features (i.e., principal components), 61 ecological traits, and 110 taxonomic traits.

**S1 Data. Predictive function of associated proteins for genes ranked by their positive and negative loading contribution to principal components 1, 4, 3 and 9.** Genes with loading values that were greater than the mean plus 1.5 times the standard deviation and less than the mean minus 1.5 times the standard deviation were included for those with positive loadings separate from those with negative loadings.

**S2 Data. Host-virus associations for link prediction based on *Orthopoxvirus* genomes extracted from NCBI.**

## References

1. Jacobs BL, Langland JO, Kibler KV, Denzler KL, White SD, Holechek SA, et al. Vaccinia virus vaccines: Past, present and future. Antiviral Research. 2009;84: 1–13. doi:10.1016/j.antiviral.2009.06.006

2. Fenner F. Global Eradication of Smallpox. Reviews of Infectious Diseases. 1982;4: 916– 930. doi:10.1093/clinids/4.5.916

3. Rimoin AW, Mulembakani PM, Johnston SC, Lloyd Smith JO, Kisalu NK, Kinkela TL, et al. Major increase in human monkeypox incidence 30 years after smallpox vaccination campaigns cease in the Democratic Republic of Congo. Proceedings of the National Academy of Sciences. 2010;107: 16262–16267. doi:10.1073/pnas.1005769107

4. Reynolds MG, Guagliardo SAJ, Nakazawa YJ, Doty JB, Mauldin MR. Understanding orthopoxvirus host range and evolution: from the enigmatic to the usual suspects. Current Opinion in Virology. 2018;28: 108–115. doi:10.1016/j.coviro.2017.11.012

5. Sutter G, Moss B. Nonreplicating vaccinia vector efficiently expresses recombinant genes. Proceedings of the National Academy of Sciences. 1992;89: 10847–10851. doi:10.1073/pnas.89.22.10847

6. Xiang Y, White A. Monkeypox virus emerges from the shadow of its more infamous cousin: family biology matters. Emerging Microbes & Infections. 2022;11: 1768–1777. doi:10.1080/22221751.2022.2095309

7. Shchelkunov SN. Orthopoxvirus genes that mediate disease virulence and host tropism. Adv Virol. 2012;2012: 524743. doi:10.1155/2012/524743

8. Shchelkunov SN. An Increasing Danger of Zoonotic Orthopoxvirus Infections. PLOS Pathogens. 2013;9: e1003756. doi:10.1371/journal.ppat.1003756

9. Mayr A, Hochstein-Mintzel V, Stickl H. Abstammung, Eigenschaften und Verwendung des attenuierten Vaccinia-Stammes MVA. Infection. 1975;3: 6–14. doi:10.1007/BF01641272

10. Rodrigues TCS, Subramaniam K, Varsani A, McFadden G, Schaefer AM, Bossart GD, et al. Genome characterization of cetaceanpox virus from a managed Indo-Pacific bottlenose dolphin (Tursiops aduncus). Virus Research. 2020;278: 197861. doi:10.1016/j.virusres.2020.197861

11. Hale VL, Dennis PM, McBride DS, Nolting JM, Madden C, Huey D, et al. SARS-CoV-2 infection in free-ranging white-tailed deer. Nature. 2022;602: 481–486. doi:10.1038/s41586-021-04353-x

12. Springer YP, Hsu CH, Werle ZR, Olson LE, Cooper MP, Castrodale LJ, et al. Novel Orthopoxvirus Infection in an Alaska Resident. Clinical Infectious Diseases. 2017;64: 1737–1741. doi:10.1093/cid/cix219

13. Bernstein AS, Ando AW, Loch-Temzelides T, Vale MM, Li BV, Li H, et al. The costs and benefits of primary prevention of zoonotic pandemics. Science Advances. 2022;8: eabl4183. doi:10.1126/sciadv.abl4183

14. Mollentze N, Babayan SA, Streicker DG. Identifying and prioritizing potential human-infecting viruses from their genome sequences. PLOS Biology. 2021;19: e3001390. doi:10.1371/journal.pbio.3001390

15. Han BA, Schmidt JP, Bowden SE, Drake JM. Rodent reservoirs of future zoonotic diseases. Proceedings of the National Academy of Sciences. 2015;112: 7039–7044. doi:10.1073/pnas.1501598112

16. Becker DJ, Albery GF, Sjodin AR, Poisot T, Bergner LM, Chen B, et al. Optimising predictive models to prioritise viral discovery in zoonotic reservoirs. The Lancet Microbe. 2022;3: e625–e637. doi:10.1016/S2666-5247(21)00245-7

17. Blagrove MS, Pilgrim J, Kotsiri A, Hui M, Baylis M, Wardeh M. Monkeypox virus shows potential to infect a diverse range of native animal species across Europe, indicating high risk of becoming endemic in the region. bioRxiv; 2022. p. 2022.08.13.503846. doi:10.1101/2022.08.13.503846

18. Fischhoff IR, Castellanos AA, Rodrigues JPGLM, Varsani A, Han BA. Predicting the zoonotic capacity of mammals to transmit SARS-CoV-2. Proc Biol Sci. 2021;288: 20211651. doi:10.1098/rspb.2021.1651

19. Wardeh M, Baylis M, Blagrove MSC. Predicting mammalian hosts in which novel coronaviruses can be generated. Nat Commun. 2021;12: 780. doi:10.1038/s41467-021-21034-5

20. Meekins DA, Morozov I, Trujillo JD, Gaudreault NN, Bold D, Carossino M, et al. Susceptibility of swine cells and domestic pigs to SARS-CoV-2. Emerging Microbes & Infections. 2020;9: 2278–2288. doi:10.1080/22221751.2020.1831405

21. Schlottau K, Rissmann M, Graaf A, Schön J, Sehl J, Wylezich C, et al. SARS-CoV-2 in fruit bats, ferrets, pigs, and chickens: an experimental transmission study. The Lancet Microbe. 2020;1: e218–e225. doi:10.1016/S2666-5247(20)30089-6

22. Haddock E, Callison J, Seifert SN, Okumura A, Tang-Huau T-L, Leventhal SS, et al. Three-Week Old Pigs Are Not Susceptible to Productive Infection with SARS-COV-2. Microorganisms. 2022;10: 407. doi:10.3390/microorganisms10020407

23. Plowright RK, Becker DJ, Crowley DE, Washburne AD, Huang T, Nameer PO, et al. Prioritizing surveillance of Nipah virus in India. PLOS Neglected Tropical Diseases. 2019;13: e0007393. doi:10.1371/journal.pntd.0007393

24. Seifert SN, Letko MC, Bushmaker T, Laing ED, Saturday G, Meade-White K, et al. Rousettus aegyptiacus Bats Do Not Support Productive Nipah Virus Replication. The Journal of Infectious Diseases. 2020;221: S407–S413. doi:10.1093/infdis/jiz429

25. Sánchez-Morales L, Sánchez-Vizcaíno JM, Pérez-Sancho M, Domínguez L, Barroso-Arévalo S. The Omicron (B.1.1.529) SARS-CoV-2 variant of concern also affects companion animals. Frontiers in Veterinary Science. 2022;9. Available: https://www.frontiersin.org/articles/10.3389/fvets.2022.940710

26. Taube JC, Rest EC, Lloyd-Smith JO, Bansal S. The global landscape of smallpox vaccination history and implications for current and future orthopoxvirus susceptibility: a modelling study. The Lancet Infectious Diseases. 2023;23: 454–462. doi:10.1016/S1473-3099(22)00664-8

27. Babayan SA, Orton RJ, Streicker DG. Predicting reservoir hosts and arthropod vectors from evolutionary signatures in RNA virus genomes. Science. 2018;362: 577–580. doi:10.1126/science.aap9072

28. Brierley L, Fowler A. Predicting the animal hosts of coronaviruses from compositional biases of spike protein and whole genome sequences through machine learning. PLOS Pathogens. 2021;17: e1009149. doi:10.1371/journal.ppat.1009149

29. Wardeh M, Blagrove MSC, Sharkey KJ, Baylis M. Divide-and-conquer: machine-learning integrates mammalian and viral traits with network features to predict virus-mammal associations. Nat Commun. 2021;12: 3954. doi:10.1038/s41467-021-24085-w

30. Gigante CM, Gao J, Tang S, McCollum AM, Wilkins K, Reynolds MG, et al. Genome of Alaskapox Virus, a Novel Orthopoxvirus Isolated from Alaska. Viruses. 2019;11. doi:10.3390/v11080708

31. Kastenmayer RJ, Maruri-Avidal L, Americo JL, Earl PL, Weisberg AS, Moss B. Elimination of A-type inclusion formation enhances cowpox virus replication in mice: Implications for orthopoxvirus evolution. Virology. 2014;452–453: 59–66. doi:10.1016/j.virol.2013.12.030

32. Fagre AC, Cohen LE, Eskew EA, Farrell M, Glennon E, Joseph MB, et al. Assessing the risk of human-to-wildlife pathogen transmission for conservation and public health. Ecology Letters. 2022;25: 1534–1549. doi:10.1111/ele.14003

33. Monkeypox – Cameroon. In: World Health Organization: Disease Outbreak News [Internet]. 5 Jun 2018 [cited 13 May 2023]. Available: https://www.who.int/emergencies/disease-outbreak-news/item/05-june-2018-monkeypox-cameroon-en

34. de Oliveira JS, Figueiredo P de O, Costa GB, de Assis FL, Drumond BP, da Fonseca FG, et al. Vaccinia Virus Natural Infections in Brazil: The Good, the Bad, and the Ugly. Viruses. 2017;9: 340. doi:10.3390/v9110340

35. Figuerola J, Green AJ. Haematozoan Parasites and Migratory Behaviour in Waterfowl. Evolutionary Ecology. 2000;14: 143–153. doi:10.1023/A:1011009419264

36. Senkevich TG, Yutin N, Wolf YI, Koonin EV, Moss B. Ancient Gene Capture and Recent Gene Loss Shape the Evolution of Orthopoxvirus-Host Interaction Genes. mBio. 12: e01495–21. doi:10.1128/mBio.01495-21

37. Carlson CJ, Gibb RJ, Albery GF, Brierley L, Connor RP, Dallas TA, et al. The Global Virome in One Network (VIRION): an atlas of vertebrate-virus associations. 2021; 2021.08.06.455442. doi:10.1101/2021.08.06.455442

38. Upham NS, Esselstyn JA, Jetz W. Inferring the mammal tree: Species-level sets of phylogenies for questions in ecology, evolution, and conservation. PLOS Biology. 2019;17: e3000494. doi:10.1371/journal.pbio.3000494

39. Soria CD, Pacifici M, Di Marco M, Stephen SM, Rondinini C. COMBINE: a coalesced mammal database of intrinsic and extrinsic traits. Ecology. 2021;102: e03344. doi:10.1002/ecy.3344

40. The IUCN Red List of Threatened Species. In: IUCN Red List of Threatened Species [Internet]. [cited 14 Jul 2023]. Available: https://www.iucnredlist.org/en

41. Faurby S, Davis M, Pedersen RØ, Schowanek SD, Antonellil A, Svenning J-C. PHYLACINE 1.2: The Phylogenetic Atlas of Mammal Macroecology. Ecology. 2018;99: 2626–2626. doi:10.1002/ecy.2443

42. Mull N, Carlson CJ, Forbes KM, Becker DJ. Virus isolation data improve host predictions for New World rodent orthohantaviruses. Journal of Animal Ecology. 2022;91: 1290–1302. doi:10.1111/1365-2656.13694

43. Elith J, Leathwick JR, Hastie T. A working guide to boosted regression trees. Journal of Animal Ecology. 2008;77: 802–813. doi:10.1111/j.1365-2656.2008.01390.x

44. De’ath G. Boosted Trees for Ecological Modeling and Prediction. Ecology. 2007;88: 243–251. doi:10.1890/0012-9658(2007)88[243:BTFEMA]2.0.CO;2

45. Benjamini Y, Hochberg Y. Controlling the False Discovery Rate: A Practical and Powerful Approach to Multiple Testing. Journal of the Royal Statistical Society: Series B (Methodological). 1995;57: 289–300. doi:10.1111/j.2517-6161.1995.tb02031.x

46. Yekutieli D, Benjamini Y. Resampling-based false discovery rate controlling multiple test procedures for correlated test statistics. Journal of Statistical Planning and Inference. 1999;82: 171–196. doi:10.1016/S0378-3758(99)00041-5

47. Washburne AD, Silverman JD, Morton JT, Becker DJ, Crowley D, Mukherjee S, et al. Phylofactorization: a graph partitioning algorithm to identify phylogenetic scales of ecological data. Ecological Monographs. 2019;89: e01353. doi:10.1002/ecm.1353

48. Holm S. A Simple Sequentially Rejective Multiple Test Procedure. Scandinavian Journal of Statistics. 1979;6: 65–70.

