## Supplemental Information for "Viral genomic features predict *orthopoxvirus* reservoir hosts"

**This PDF file includes:**

Supporting text

Figures S1 to S9

Tables S1 to S7

Legends for Datasets S1 to S2

SI References

**Other supporting materials for this manuscript include the following:**

Datasets S1 to S2

Supporting Information Text

**Supporting Methods**

***Host-virus association data***

We obtained known host-OPV associations from the Global Virome, in One Network (<https://www.viralemergence.org/virion>; VIRION), an open database of host-virus interactions drawn from scientific literature and online databases (<https://github.com/viralemergence/virion>). As one of the most comprehensive, published databases on the mammal virome, VIRION consists of over 23,000 unique interactions representing 9,521 virus species and 3,692 vertebrate host species reconciled to NCBI taxonomy. We filtered the dataset to include only interactions with OPVs that were detected via PCR or virus isolation. Due to potential inaccuracies in viral detection, interactions with variola virus were excluded from analysis as the virus was eradicated prior to the development of many modern diagnostics. We then collapsed associations to the genus level and kept only unique interactions, resulting in a total of 95 unique host-virus associations between 71 mammal genera and 19 virus “species.” To allow for out-of-sample predictions of host genera with unknown host-virus interactions, we merged host positivity data with the broader mammal taxonomy [1]. We then assigned pseudoabsences to mammal genera from taxonomic orders in which known host-virus associations exist, resulting in an additional 875 mammal genera labeled as pseudo absences. Our final dataset of mammal genera consisted of 946 mammal genera including 71 with known OPV positivity: 58 were detected via PCR and 48 were detected via virus isolation (28 were OPV positive by both PCR and virus isolation).

For the link prediction model, we merged data from VIRION with host-OPV associations obtained from the National Center for Biotechnology Information (NCBI). In total, 197 orthopoxvirus genomes with annotations, when available, were downloaded from NCBI, representing 45 unique host-virus associations between 35 host genera and 12 virus species. In addition, we identified host-virus associations from experimental studies when the specific OPV isolate had been included (Dataset S1). After merging sequence data with data from VIRION, the final known association dataset for the link prediction model described a network of 254 host-virus associations between 75 host genera and 19 virus species. Next, we merged host-virus association data with the broader mammal taxonomy [1] to accommodate out-of-sample predictions of host genera with unknown host-virus interactions. Keeping only those mammal genera that exist in taxonomic orders with known host-virus associations, we expanded our mammal-virus network to include all combinations of mammal genera and OPVs. We then assigned pseudo absences to those host-virus associations with no known OPV detection (by PCR or virus isolation), resulting in an additional 17,873 pseudo-absent links. The final mammal-virus network was composed of 18,126 host-virus pairs between 946 distinct mammal genera and 19 OPV species.

***Viral predictor data pre-processing***

To incorporate viral genomic features in our link prediction model, we extracted genomes and their annotations from NCBI for all OPVs and aligned them using MAFFT v 7.490 [2,3]. We identified accessory genes using Roary implemented in Galaxy v 3.13.0 + galaxy2 with 80% minimum sequence identity for blastp and 95% of isolates a gene must be in to be designated as part of the core genome. The resulting 981 accessory genes were transformed into a matrix of binary presence/absence variables for the 197 sequence-based host-OPV associations.

Next, we conducted principal components analysis (PCA) to distill the 981 accessory gene variables down to their most important features for incorporation in the link prediction model. PCA linearly transforms high dimensional datasets into a coordinate system of uncorrelated axes where the first principal component accounts for the greatest amount of variance, followed by each subsequent component thereafter. Thus, PCA captures most of the variance in the data in as a few dimensions as possible. Using the R package *stats*, we conducted four PCAs: the first included all 981 variables and 197 sequences; the second excluded accessory gene variables with close to zero variance (i.e., accessory genes present in only one virus species); the third excluded sequences with identical patterns of accessory gene presence/absence as another sequence in our dataset; and the fourth excluded potential outlying scores (sequences).

Each PCA produced marginal differences in the proportion of variance explained by each dimension. Furthermore, we observed no noticeable differences in the spatial distribution of scores (i.e., the coordinates of each individual sequence) and loadings (i.e., the correlations between each original variable and the principal components). As such, we used results from our first PCA and extracted the scores of the first 10 PCs as our final list of viral predictor variables. These ten PCs explained roughly 70% of the variance in the data (Figure S4). Because data for the ten PC variables were only available for the host-virus associations obtained from sequence data for 12 OPV species, we conducted median imputation for associations with any of the 12 OPV species for which PC data were missing.

To identify how OPV sequences group in our coordinate system of principal components, we conducted hierarchical clustering analysis (HCA) on our ten principal components using the *clustree package* in R. In HCA, each sequence is initially assigned its own cluster and then at each step of the algorithm, the two most similar clusters are joined until a single cluster is left, forming a bottom-up dendogram. As the number of clusters (*k*) increased, we examined how sequences changed groupings to determine which clusters were distinct and which were unstable, and we calculated the within-cluster sum of squared errors (intra-cluster variance) to help determine the optimal number of clusters (i.e., the elbow method). Results of hierarchical clustering can be found on the GitHub repository.

***Phylogeny***

To capture mammal phylogeny, we used a supertree of extant mammal species [1] trimmed to the genus level using the *treespace* package in R. We also reconstructed the viral phylogeny of the 12 OPV species represented in the 197 sequences/associations extracted from GenBank to investigate clustering in the distribution of viral genomic data. Multiple sequence alignment of viral genomes was completed using MAFFT v7.490 and a maximum likelihood tree was inferred with PhyML v3.3.20180621 using a generalized time reversible substitution model and estimating the gamma distribution parameter and proportion of invariable sites.

***Taxonomic reconciliation***

We corrected naming inconsistencies by manually matching host trait dataset names to those of the mammal phylogeny. This included reverting names to their homotypic synonyms: *Liomys* to *Heteromys*, *Oreonax* to *Lagothrix*, *Paralomys* to *Phyllotis*, *Pearsonomys* to *Geoxus*, *Pipanacoctomys* to *Tympanoctomys*, and *Pseudalopex* to *Lycalopex* [4,5]. Additionally, the genus name *Classomys* in our host trait dataset was missing from the mammal supertree and was switched to *Delomys*, a phylogenetically closely related genus of the same *Sigmodontinae* superfamily and a sister taxa with many morphological similarities [6].

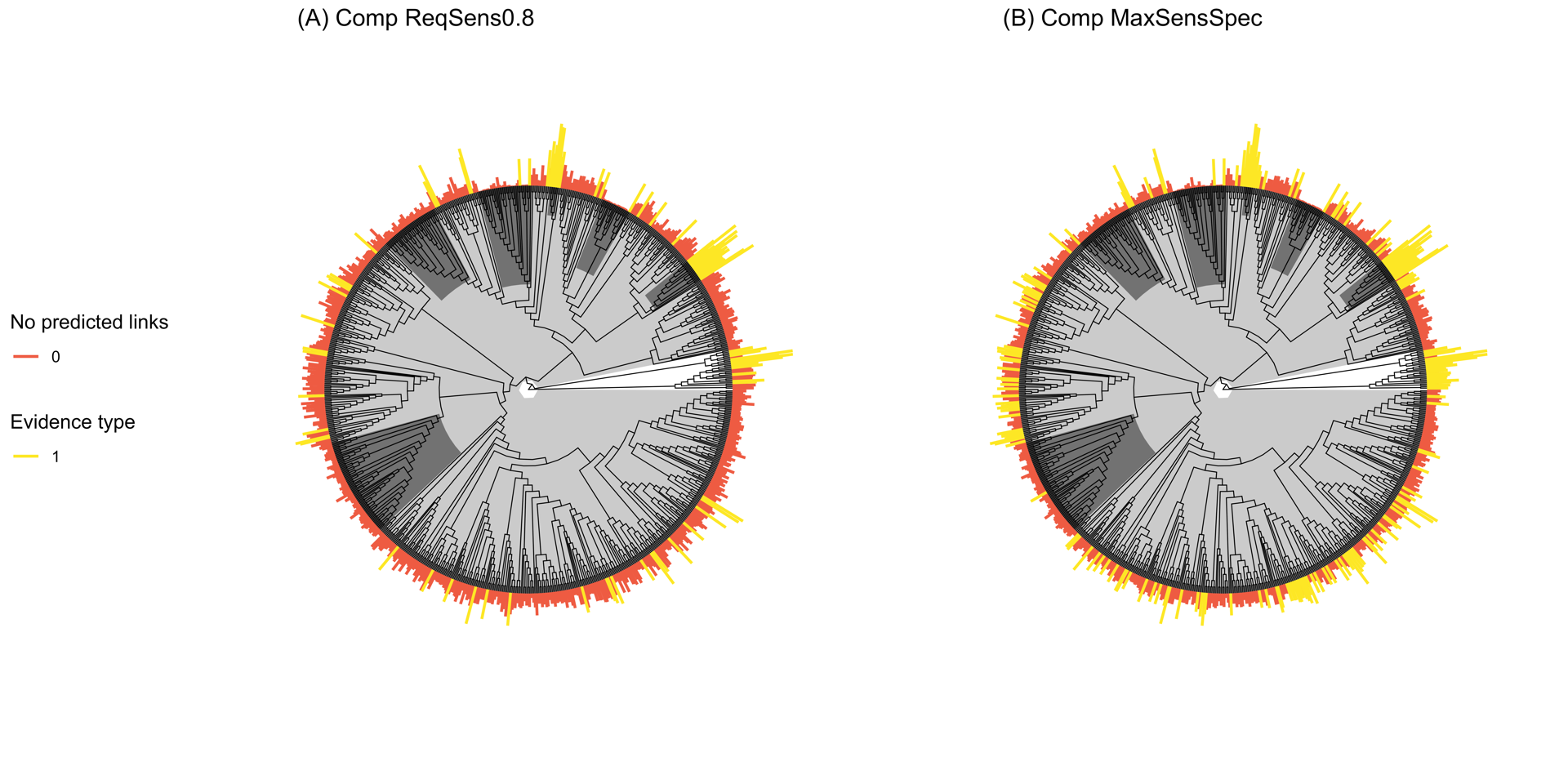

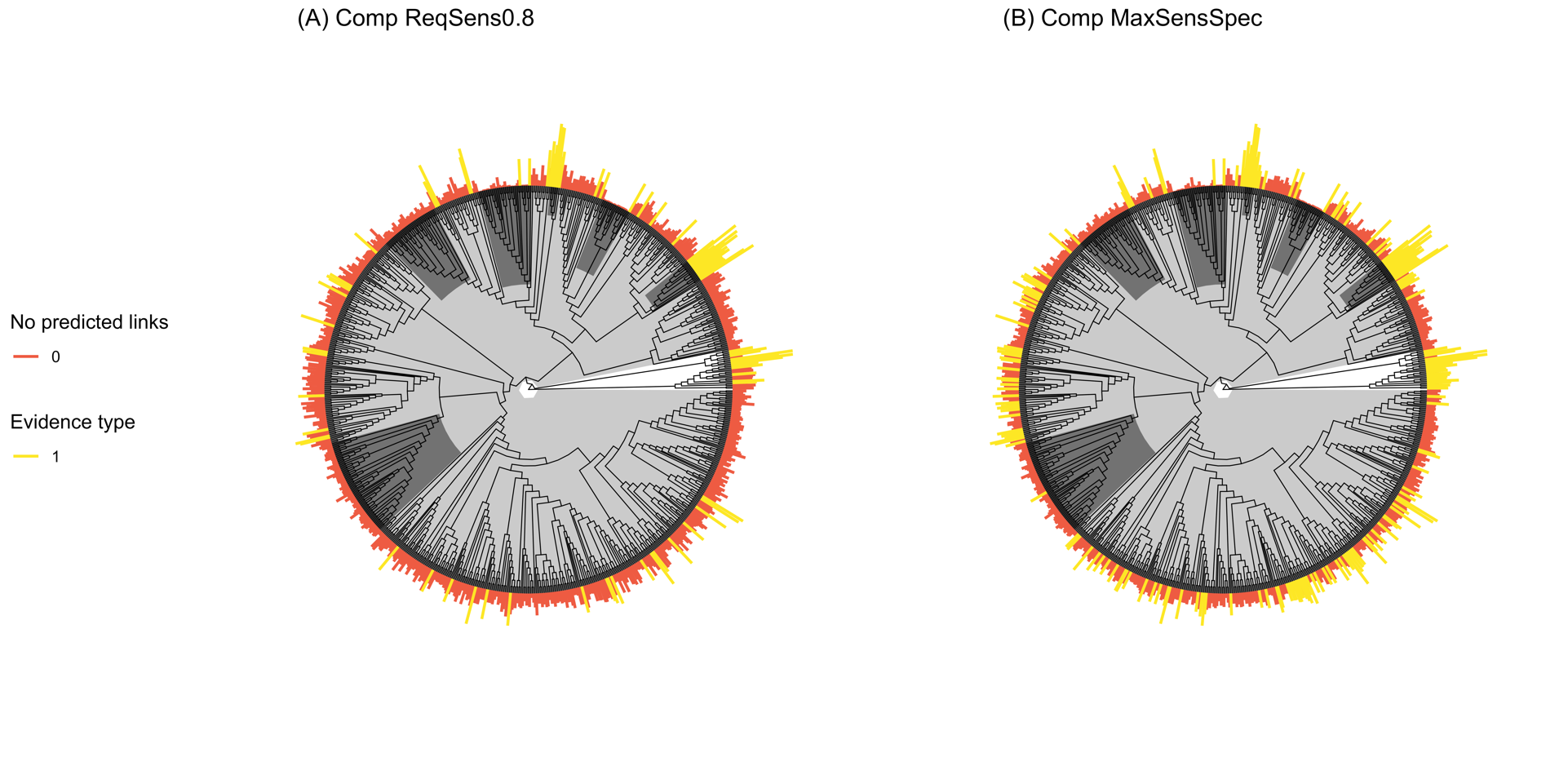

**Figure S1. Predicted probabilities of orthopoxvirus positivity and taxonomic patterns in predictions identified through phylogenetic factorization** **for the host prediction model trained on virus isolation data as the response (i.e, susceptible host model).** Segments are scaled by probabilities and colored yellow for those host genera predicted to be potential reservoirs based on **(A)** an 80% sensitivity threshold versus **(B)** a threshold maximizing the sum of sensitivity and specificity. Clades identified through phylogenetic factorization with significantly different mean predictions are shaded in grey.

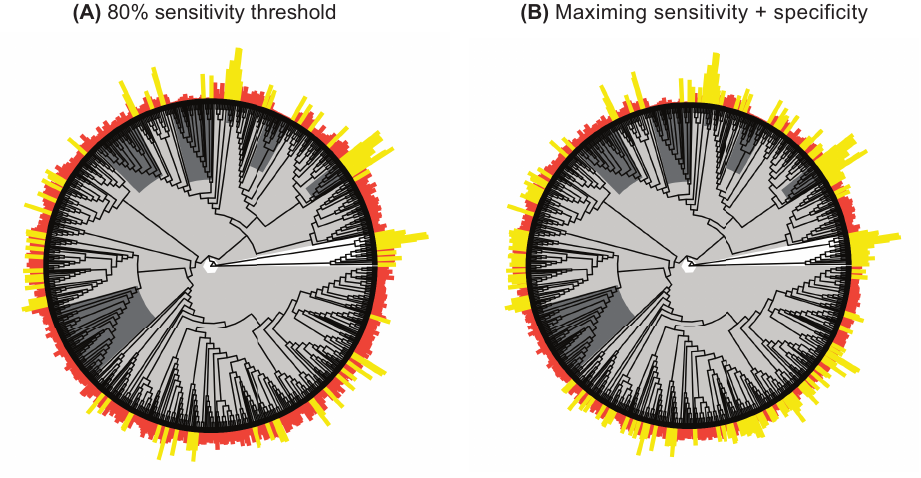

Figure S2. Trait profiles of mammal genera for the host prediction models trained on (A) PCR data (i.e., host exposure model) and (B) virus isolation data (i.e., susceptible host model) as the response variable. Partial dependence plots for the top ten predictors across BRTs applied to 100 random partitions of training and test data are displayed in order of relative importance. Grey lines or points show the marginal effect of a given variable for prediction of host status from each data partition; black lines show the average marginal effect. Histograms and rug plots display the distribution of continuous and categorical predictor variables, respectively, across all included mammal genera.

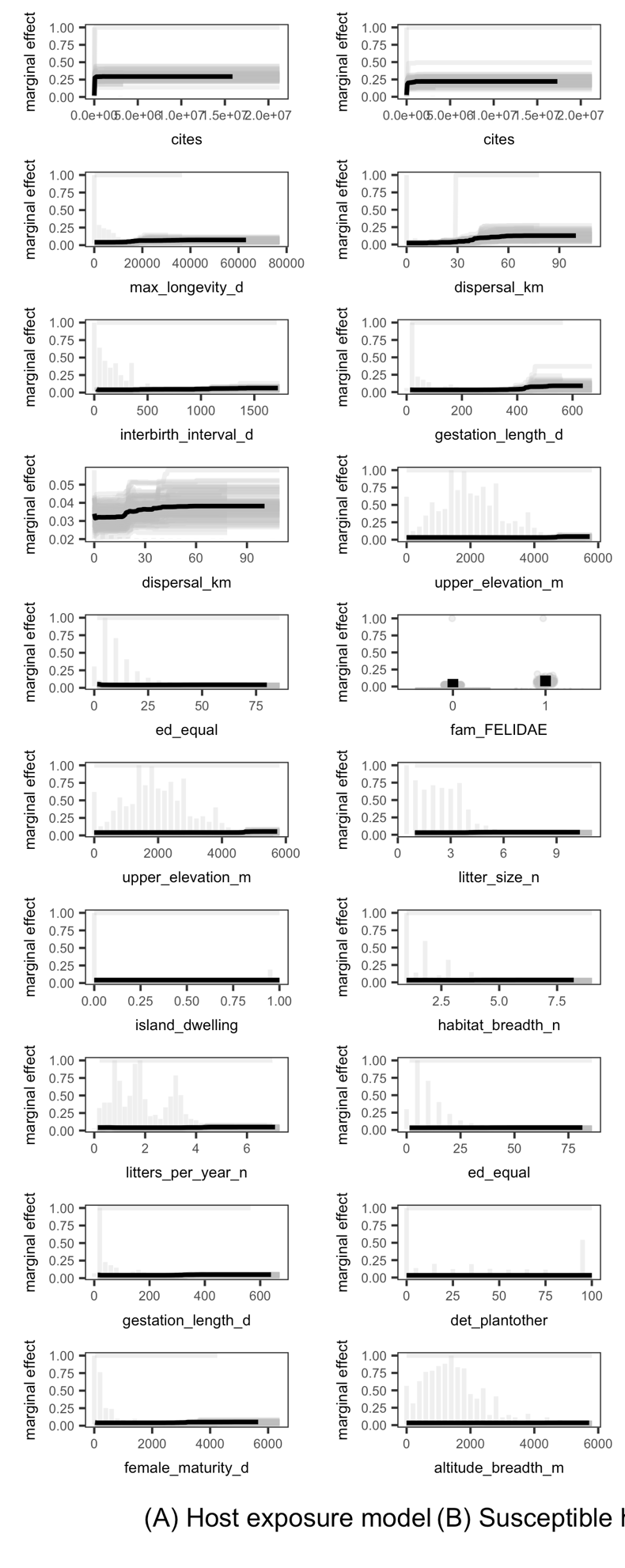

**(A)** Host exposure model	**(B)** Susceptible host model

Figure S3. Trait profiles of mammal genera for the link prediction model when trained on (A) host and viral features versus (B) host traits only. Partial dependence plots for the top ten predictors across BRTs applied to 100 random partitions of training and test data are displayed in order of relative importance. Grey lines or points show the marginal effect of a given variable for prediction of host status from each data partition. Histograms and rug plots display the distribution of continuous and categorical predictor variables, respectively, across all included mammal genera.

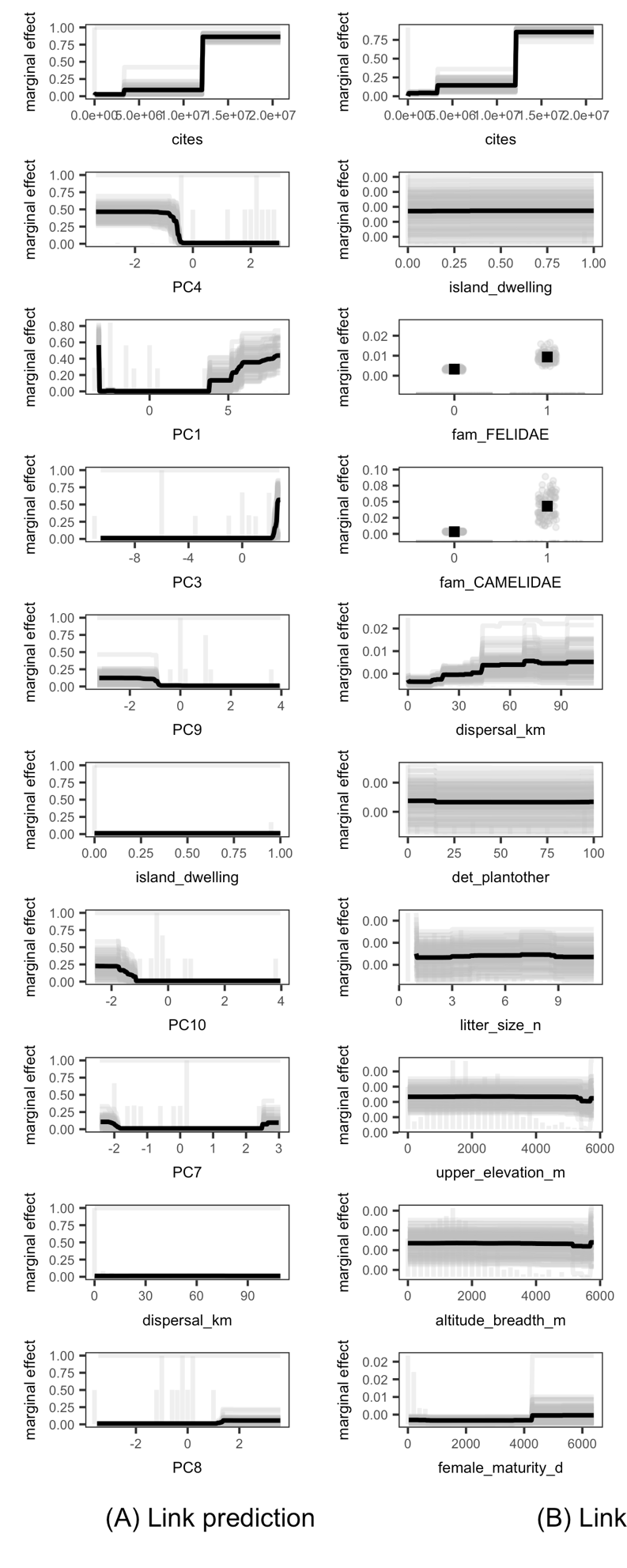

**(A)** Link prediction model

**(B)** Link prediction model trained only on host traits

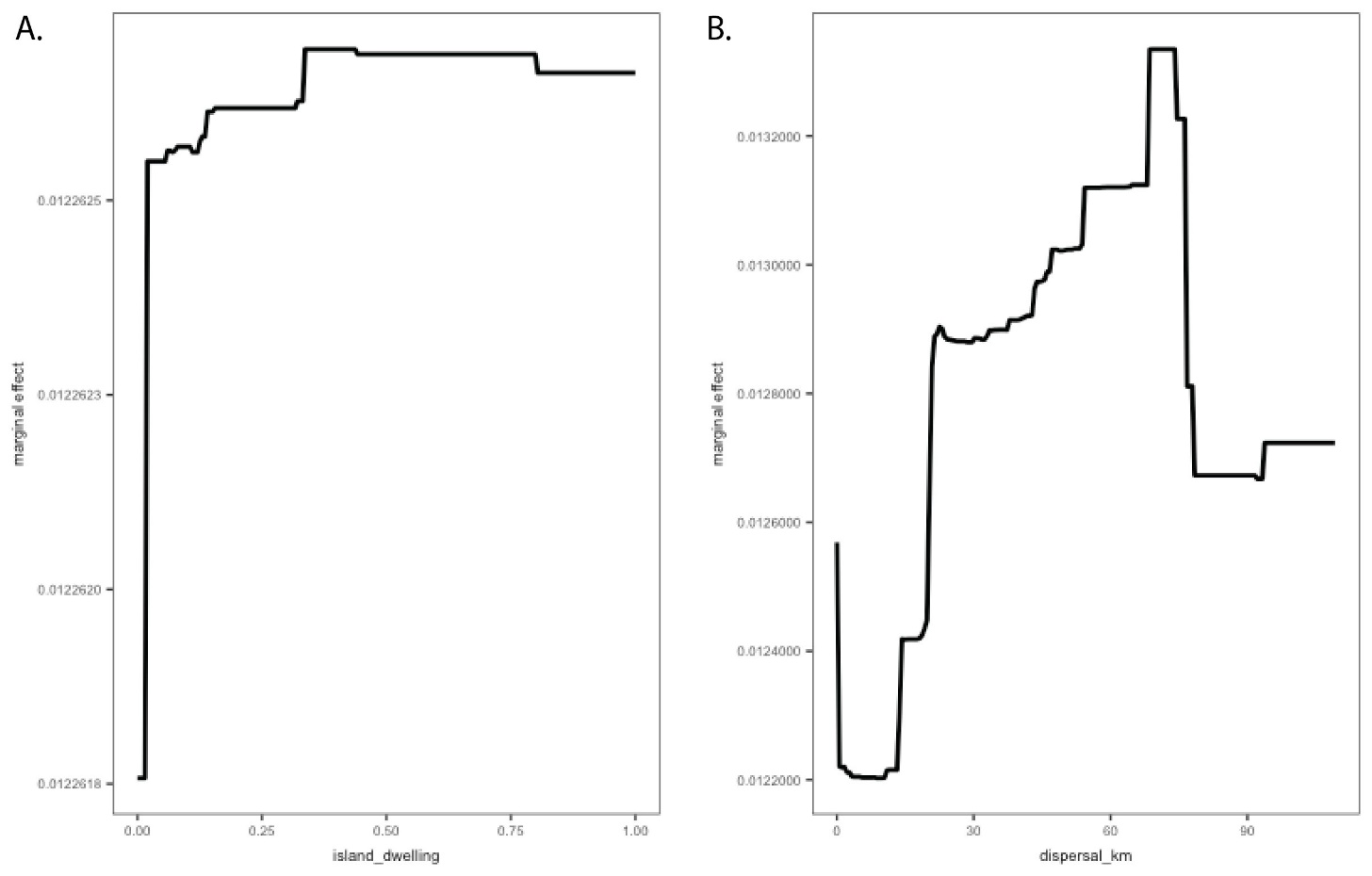

Figure S4. Marginal effects of the host traits with highest relative importance in the link prediction model when trained on host and viral features. Partial dependence plots for island dwelling (A) and dispersal (B) across BRTs applied to 100 random partitions of training and test data are displayed in order of relative importance.

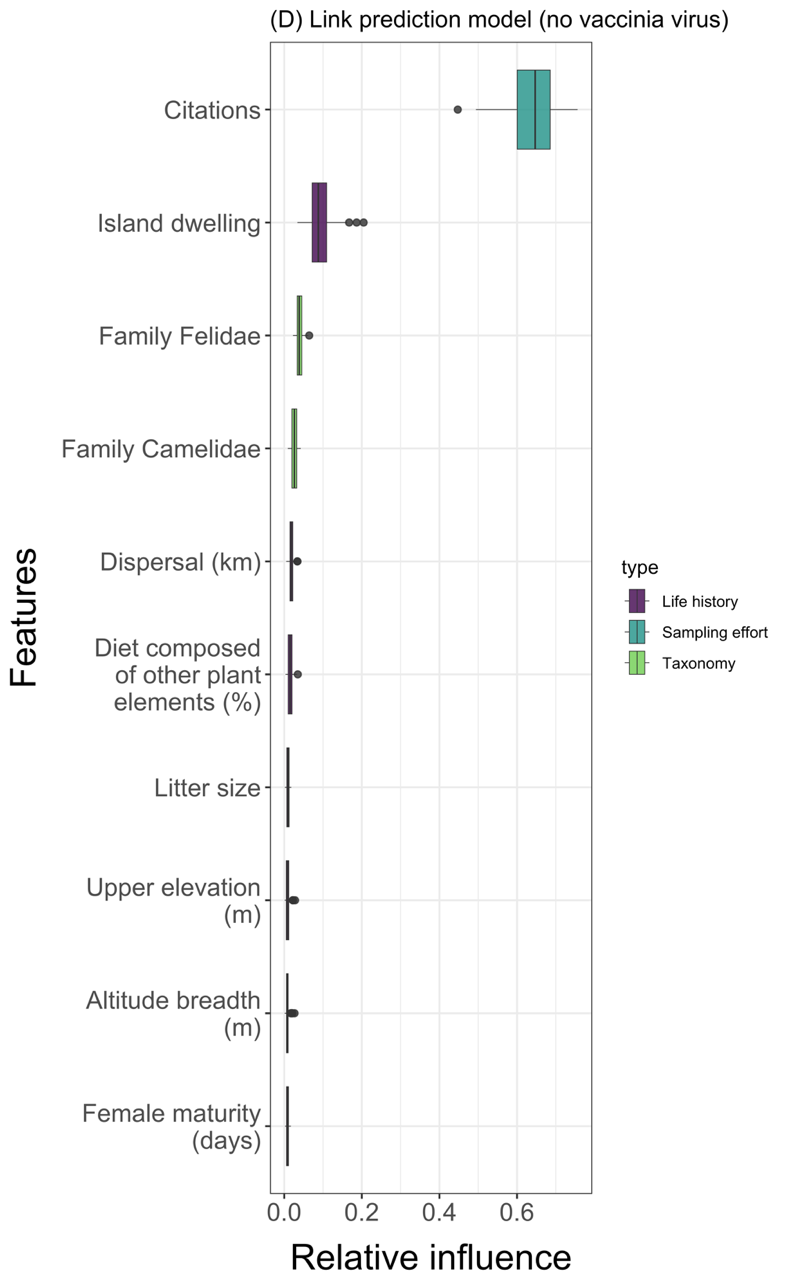

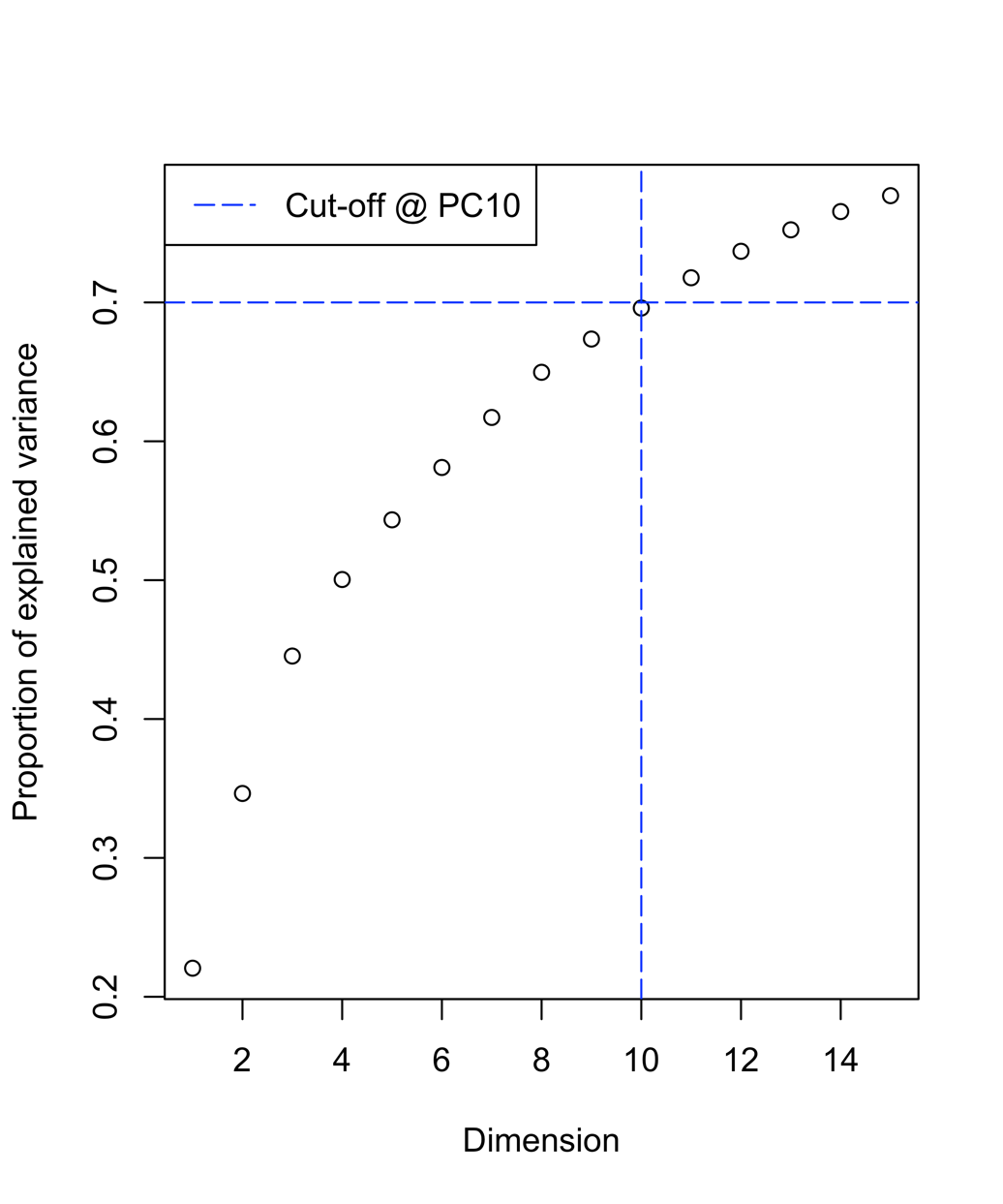

**Figure S5. Cumulative variance plot for the first ten principal components (PC).**

**Figure S6. Relative influence of model features ranked for the link prediction model trained only on host traits.** Each horizontal bar plots the variability in the relative influence of the predictor variables as measured across 100 random partitions of training (70%) and test (30%) data. Boxplots show the median and interquartile range, whiskers are the extremes, and circles are additional outliers.

**Figure S7. Plot of orthopoxvirus accessory gene loadings by the predicted gene function for principal components (A) 1 and 2, (B) 3 and 4, (C) 5 and 6, (D) 7 and 8, and (E) 9 and 10.** Only genes with loading values that were greater than or less than 1.5 times the standard deviation from the mean are represented. Genes predicted to play a role in host interaction and immune evasion include those predicted to encode the Ankyrin family of proteins, Kelch-like proteins, and proteins involved in chemokine/cytokine regulation, cell death/cell cycle regulation, pathogen recognition, and antagonizing host adaptive immunity.

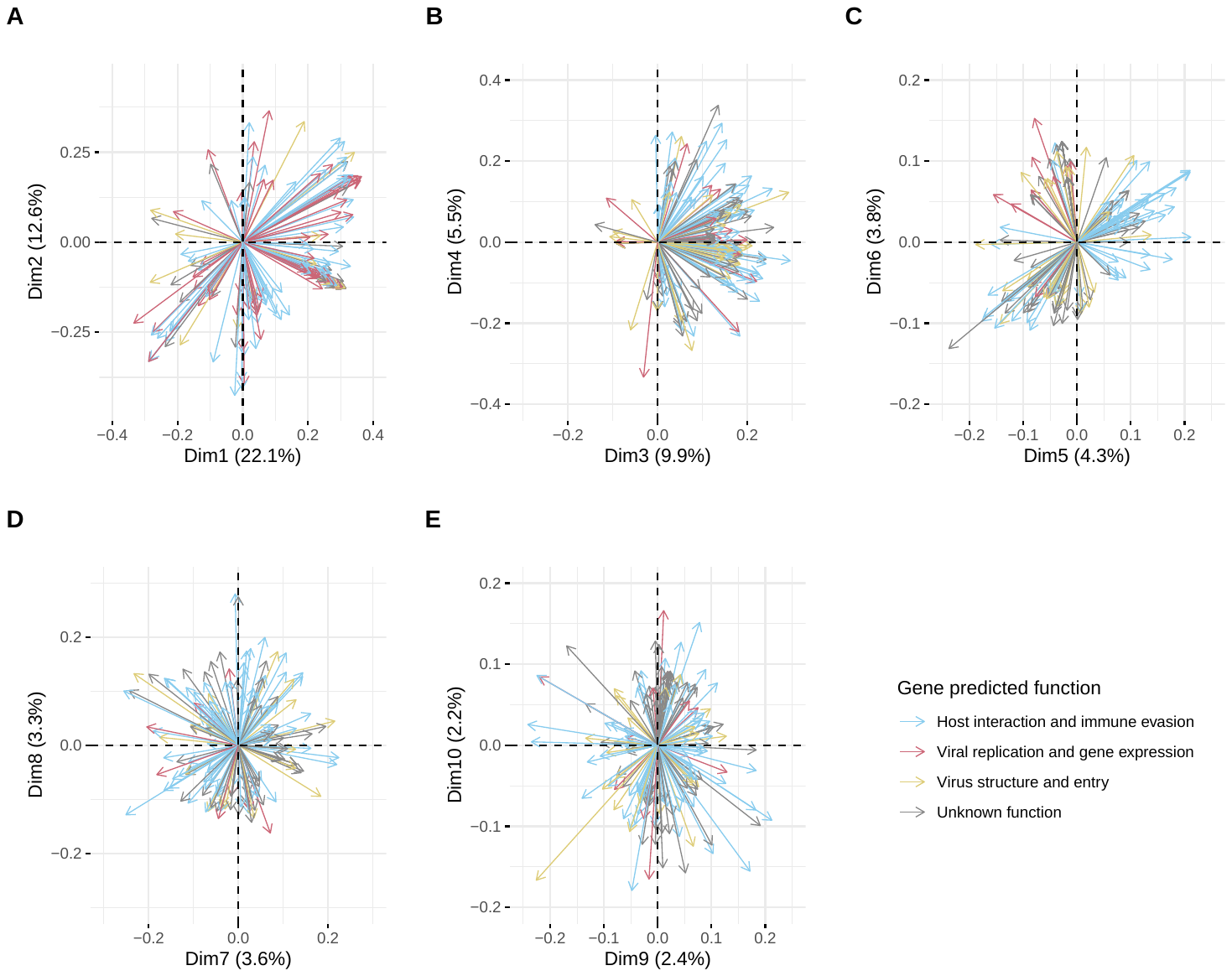

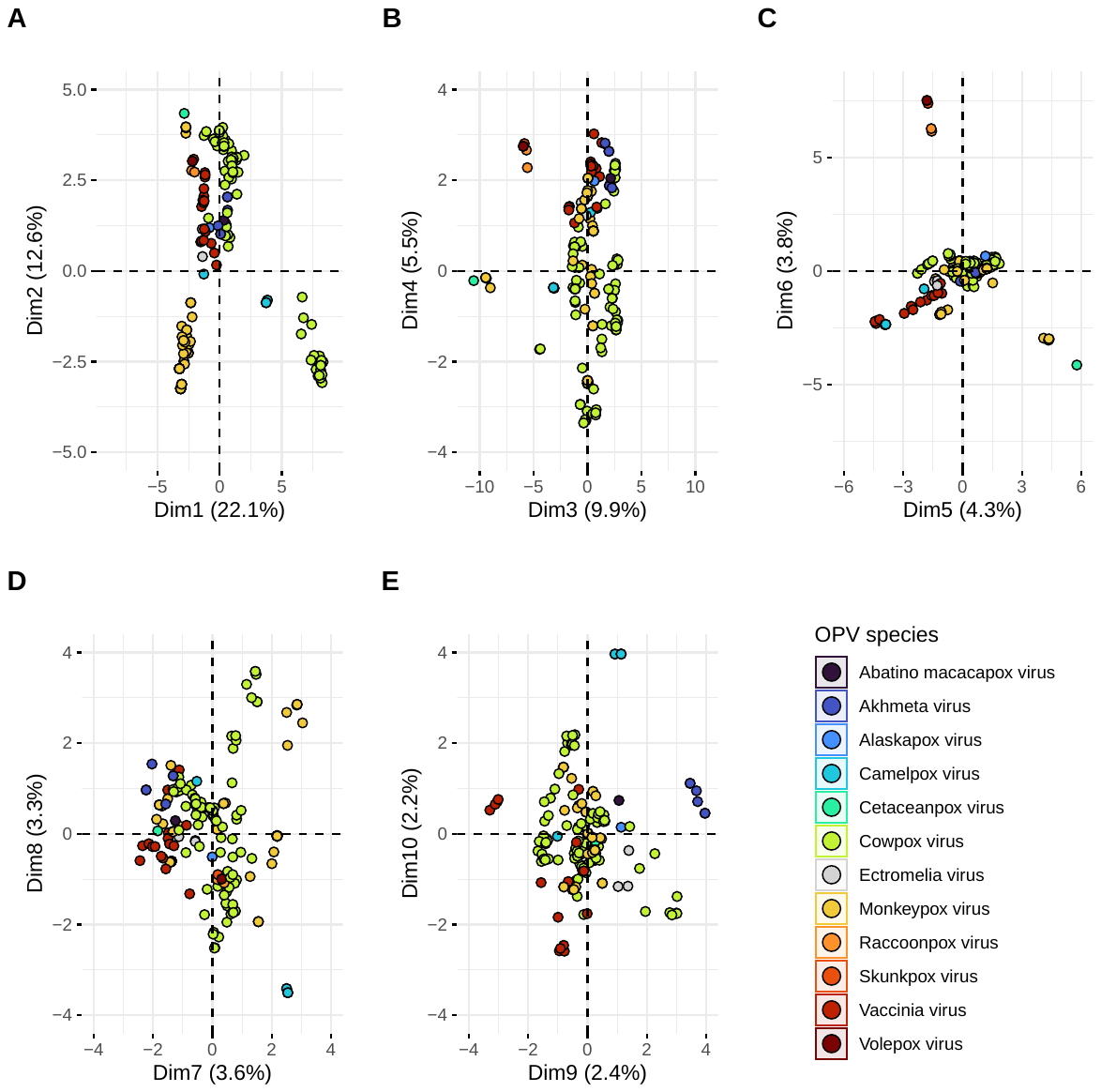

**Figure S8. Plot of orthopoxvirus sequences by viral species for principal components (A) 1 and 2, (B) 3 and 4, (C) 5 and 6, (D) 7 and 8, and (E) 9 and 10.**

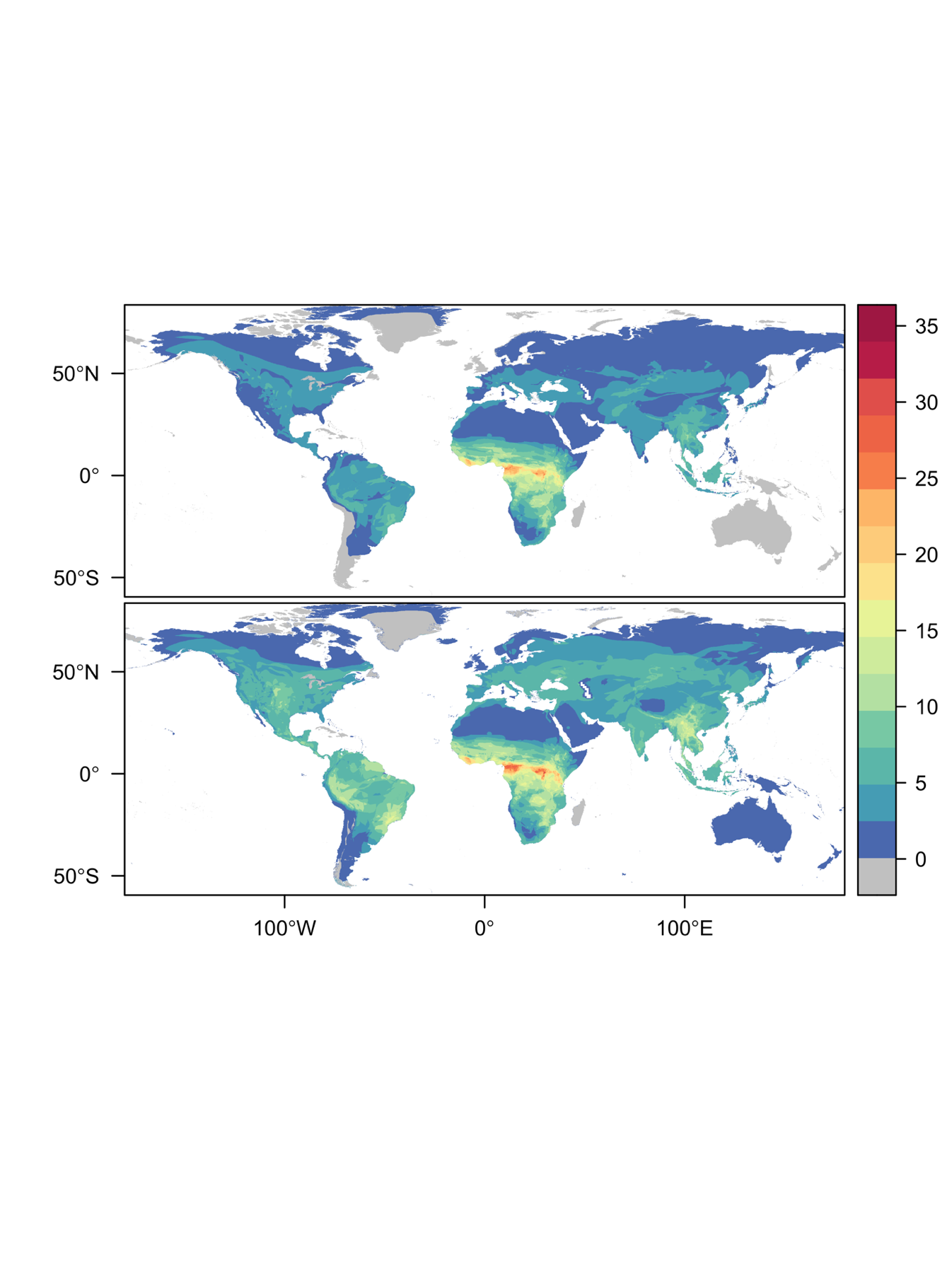

**(A)**

Observed hosts

Predicted hosts

**(B)**

**Figure S9. Geographic distribution of mpox virus hosts.** Host distributions are based on the IUCN Red List database of mammal geographic ranges for those species belonging to (A) observed host genera and (B) predicted (both observed and unobserved) host genera based on the results of the link prediction model and applying a 90% sensitivity threshold for host classification. The corresponding legend depicts the number of species with overlapping geographic range by color.

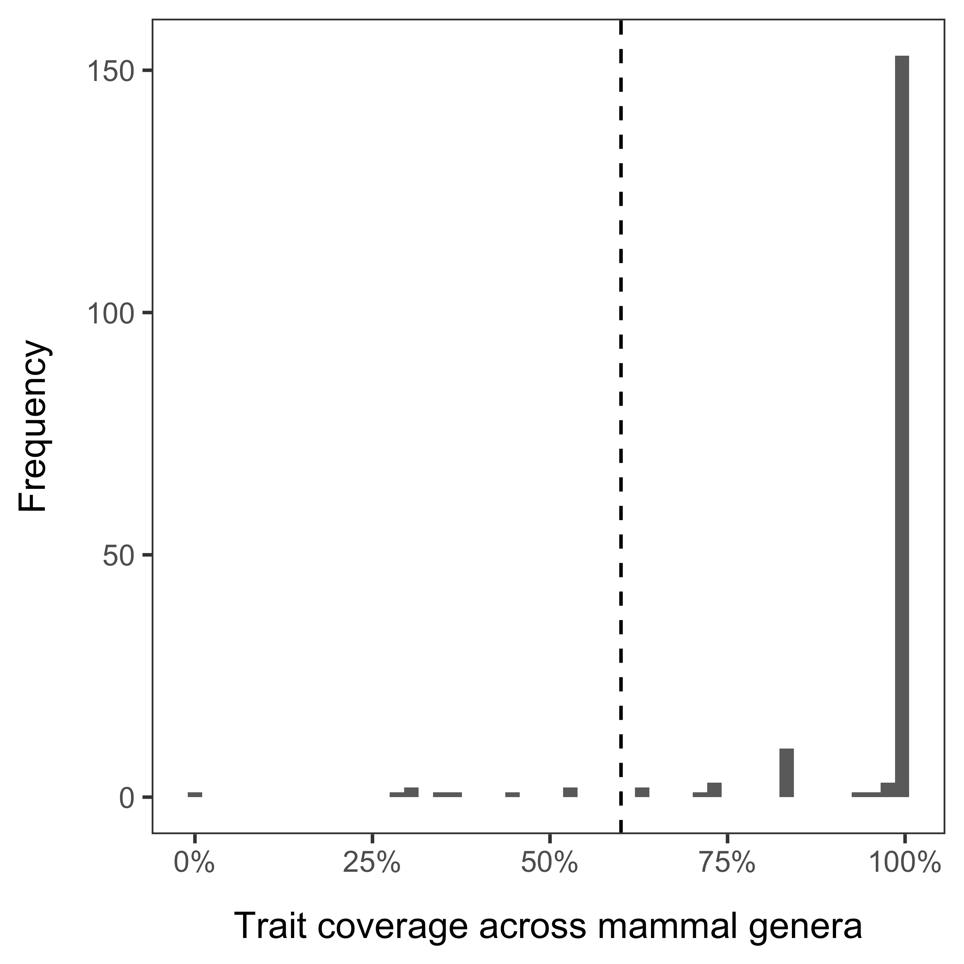

**Figure S10. Host trait coverage across mammal genera.** Features with at least 60% coverage (denoted by the dashed line) across mammal genera were included in the BRT models. A complete list of feature coverage is available on the GitHub repository.

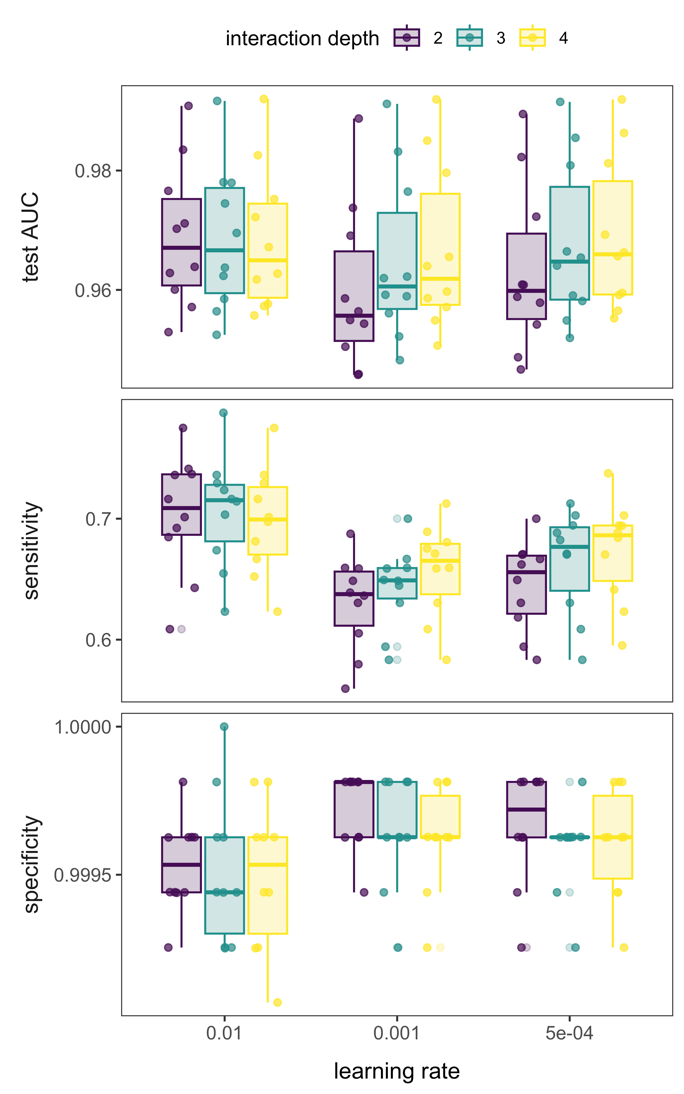

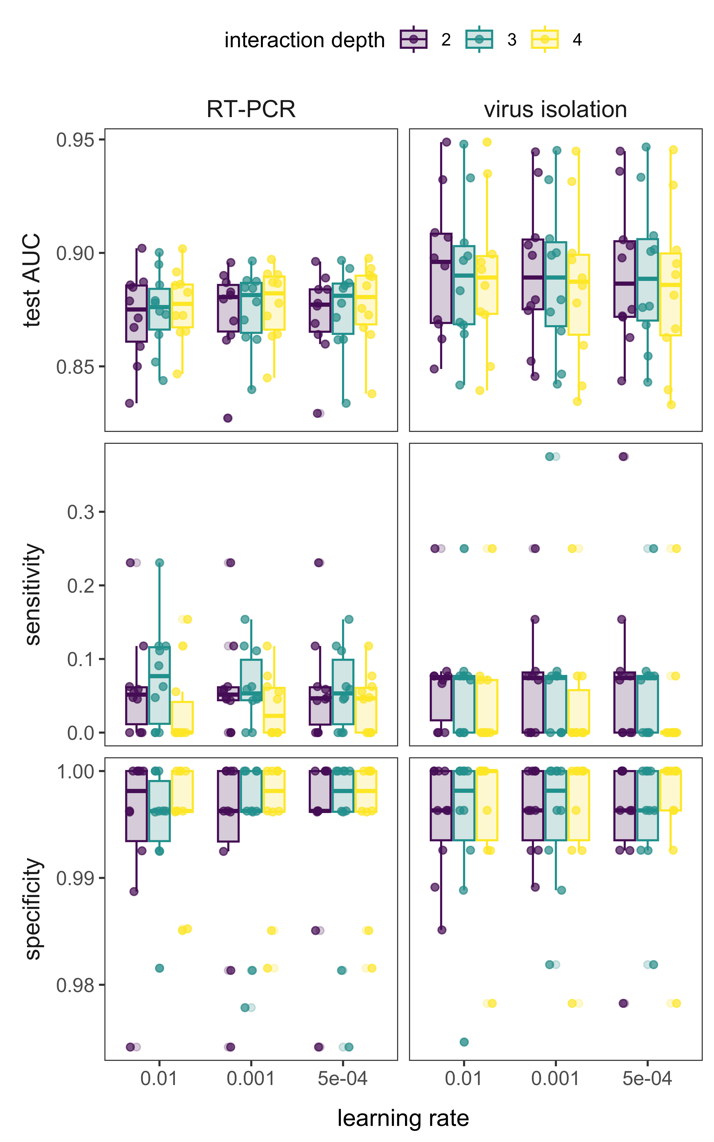

(B) Virus isolation

**(C)** Link prediction

(A) RT-PCR

**Figure S11. Performance measures for host trait BRT models trained on (A) RT-PCR versus (B) virus isolation data as the response and (C) link prediction BRT models during parameter tuning.** Boxplots show the median and interquartile range alongside raw data for all 10 random splits of training (70%) and test (30%) data for each combination of learning rate and interaction depth.

| **Table S1. Mean performance measures for each ensemble model and their corresponding standard errors (SE) were calculated based on the area under the receiver operating characteristic curve (AUC), the sensitivity, and the specificity for each model assuming a threshold value of 0.5**. We used 100 random partitions to generate an ensemble. An unpaired *t*-test with *p*-values adjusted for the false discovery rate using the Benjamini Hochberg correction compares the performance of the host exposure model (based on molecular detection of viral DNA using PCR techniques) to the susceptible host model (based on virus isolation from hosts) and that of the link prediction models with vs. without vaccinia virus associations. The effect size for the *t*-test was calculated using Cohen's *d,* which standardizes the mean difference. | | | | |
| --- | --- | --- | --- | --- |
|  | AUC | Sensitivity | | Specificity |
|  | (mean ± SE) | (mean ± SE) | | (mean ± SE) |
| Host exposure model | 0.86 ± 0.003 | 0.46 ± 0.013 | | 0.96 ± 0.003 |
| Susceptible host model | 0.88 ± 0.003 | 0.35 ± 0.014 | | 0.99 ± 0.001 |
| *t* | -4.38** | 5.66** | | -7.21** |
| Cohen's *d* | -0.62 | 0.8 | | -1.02 |
| Link prediction model | 0.96 ± 0.000 | 0.70 ± 0.003 | | 9.98 x 10^-1^ ± 0.000 |
| Link prediction model excluding vaccinia virus associations | 0.95 ± 0.001 | 0.74 ± 0.003 | | 9.97 x 10^-1^± 0.000 |
| *t* | 10.1** | -10.06** | | 2.99* |
| Cohen's *d* | 1.43 | -1.42 | | 0.42 |
| Link prediction model trained on host traits only | 0.75 ± 0.006 | 0.15 ± 0.004 | | 0.99 ± 0.000 |
| * significant at *p* < 0.01; ** significant at *p* < 0.001 | | |  | |

| **Table S2. Phylogenetic factorization of mean predicted probabilities for orthopoxvirus positivity for the host exposure model** (**based on molecular detection of viral DNA using PCR techniques).** The number of retained clades after a 5% family-wise error rate, taxa corresponding to those clades, number of species per clade, and mean predicted probabilities for the clade compared to the paraphyletic remainder are shown. | | | | | |
| --- | --- | --- | --- | --- | --- |
| Factor | Taxa | Tips | Clade | <Other< | Clade |
| 1 | Lagomorpha, Rodentia | 514 | 0.22 | 0.29 | -- |
| 2 | Phocoenidae, Monodontidae, Orcinus, Orcaella, Grampus, Pseudorca, Peponocephala, Globicephala, Feresa, Steno, Sotalia, Tursiops, Lagenodelphis, Sousa, Stenella, Delphinus, Lagenorhynchus, Cephalorhynchus | 21 | -- | 0.24 | 0.45 |
| 3 | Hylobatidae, Hominidae, Cercopithecidae | 29 | -- | 0.24 | 0.42 |
| 4 | Felidae | 17 | -- | 0.25 | 0.43 |
| 5 | Diatomyidae, Ctenodactylidae, Hystricidae, Thryonomyidae, Petromuridae, Heterocephalidae, Bathyergidae, Cuniculidae, Dasyproctidae, Caviidae, Erethizontidae, Dinomyidae, Chinchillidae, Capromyidae, Echimyidae, Myocastoridae, Ctenomyidae, Octodontidae, Abrocomidae | 74 | 0.16 | 0.26 | -- |
| 6 | Perissodactyla | 8 | -- | 0.25 | 0.43 |

| **Table S3. Phylogenetic factorization of mean predicted probabilities for orthopoxvirus positivity for the susceptible host model (based on virus isolation from hosts).** The number of retained clades after a 5% family-wise error rate, taxa corresponding to those clades, number of species per clade, and mean predicted probabilities for the clade compared to the paraphyletic remainder are shown. | | | | | |
| --- | --- | --- | --- | --- | --- |
| Factor | Taxa | Tips | Clade | <Other< | Clade |
| 1 | Felidae | 17 | -- | 0.16 | 0.47 |
| 2 | Tapiridae, Rhinocerotidae | 6 | -- | 0.16 | 0.55 |
| 3 | Eulipotyphla, Carnivora, Perissodactyla, Cetartiodactyla, Primates, Lagomorpha, Rodentia | 894 | 0.15 | 0.39 | -- |
| 4 | Ziphiidae, Physeteridae, Platanistidae, Iniidae, Phocoenidae, Monodontidae, Delphinidae, Neobalaenidae, Balaenopteridae, Eschrichtiidae, Balaenidae | 40 | 0.06 | 0.17 | -- |
| 5 | Phocidae, Odobenidae, Otariidae | 22 | 0.05 | 0.17 | -- |
| 6 | Diatomyidae, Ctenodactylidae, Hystricidae, Thryonomyidae, Petromuridae, Heterocephalidae, Bathyergidae, Cuniculidae, Dasyproctidae, Caviidae, Erethizontidae, Dinomyidae, Chinchillidae, Capromyidae, Echimyidae, Myocastoridae, Ctenomyidae, Octodontidae, Abrocomidae | 74 | 0.11 | 0.17 | -- |
| 7 | Nesotragus, Neotragus, Aepyceros, Pelea, Redunca, Kobus, Procapra, Ourebia, Raphicerus, Madoqua, Dorcatragus, Saiga, Nanger, Eudorcas, Gazella, Antilope, Litocranius, Antidorcas, Oreotragus, Philantomba, Sylvicapra, Cephalophus, Pantholops, Ovibos, Naemorhedus, Capricornis, Ovis, Nilgiritragus, Hemitragus, Capra, Pseudois, Budorcas, Myotragus, Oreamnos, Rupicapra, Arabitragus, Ammotragus, Connochaetes, Damaliscus, Beatragus, Alcelaphus, Hippotragus, Oryx, Addax | 44 | 0.1 | 0.17 | -- |

| **Table S4. Phylogenetic factorization of mean predicted probabilities for host-orthopoxvirus associations for the link prediction model trained on host and viral features.** The number of retained clades after a 5% family-wise error rate, taxa corresponding to those clades, number of species per clade, and mean predicted probabilities for the clade compared to the paraphyletic remainder are shown. | | | | | |
| --- | --- | --- | --- | --- | --- |
| Factor | Taxa | Tips | Clade | <Other< | Clade |
| 1 | Felidae | 17 | -- | 0.02 | 0.08 |
| 2 | Lagomorpha, Rodentia | 514 | 0.02 | 0.03 | -- |
| 3 | Perissodactyla | 8 | -- | 0.02 | 0.06 |
| 4 | Physeteridae, Platanistidae, Berardius, Mesoplodon, Hyperoodon, Ziphius, Tasmacetus, Iniidae, Phocoenidae, Monodontidae, Delphinidae, Neobalaenidae, Balaenopteridae, Eschrichtiidae, Balaenidae | 39 | -- | 0.02 | 0.04 |

| **Table S5. Estimation of threshold values (*t*) based on different optimal thresholding methods for the host exposure model (based on molecular detection of viral genomes using PCR techniques), the susceptible host model (based on virus isolation from hosts), and the link prediction model trained on host and viral features.** Changes in the number of predicted hosts (*n*), sensitivity, and specificity were calculated for each threshold value applied to each ensemble model. | | | | | |
| --- | --- | --- | --- | --- | --- |
|  | Method^1^ | *t* | *n* | Sensitivity | Specificity |
| Host exposure model | Sens=Spec^2^ | 0.321 | 172 | 0.862 | 0.863 |
|  | MaxSensSpec^3^ | 0.355 | 126 | 0.828 | 0.912 |
|  | MaxKappa^4^ | 0.451 | 60 | 0.655 | 0.975 |
|  | MaxPCC^5^ | 0.507 | 32 | 0.414 | 0.991 |
|  | ReqSens^6^ 80% | 0.361 | 116 | 0.810 | 0.922 |
|  | ReqSens 85% | 0.328 | 166 | 0.862 | 0.869 |
|  | ReqSens 90% | 0.284 | 263 | 0.914 | 0.764 |
|  | ReqSens 95% | 0.214 | 520 | 0.966 | 0.477 |
| Susceptible host model | Sens=Spec | 0.234 | 152 | 0.878 | 0.872 |
|  | MaxSensSpec^2^ | 0.217 | 185 | 0.951 | 0.839 |
|  | MaxKappa^3^ | 0.321 | 61 | 0.756 | 0.967 |
|  | MaxPCC^4^ | 0.404 | 31 | 0.512 | 0.989 |
|  | ReqSens^5^ 80% | 0.287 | 82 | 0.805 | 0.946 |
|  | ReqSens 85% | 0.254 | 120 | 0.854 | 0.906 |
|  | ReqSens 90% | 0.228 | 164 | 0.902 | 0.860 |
|  | ReqSens 95% | 0.217 | 185 | 0.951 | 0.839 |
| Link prediction model | Sens=Spec | 0.044 | 1864 | 0.901 | 0.901 |
|  | MaxSens+Spec | 0.057 | 1448 | 0.891 | 0.924 |
|  | MaxKappa^3^ | 0.411 | 104 | 0.604 | 0.998 |
|  | MaxPCC^4^ | 0.632 | 52 | 0.416 | 0.999 |
|  | ReqSens^5^ 80% | 0.128 | 502 | 0.802 | 0.976 |
|  | ReqSens 85% | 0.070 | 1235 | 0.851 | 0.936 |
|  | ReqSens 90% | 0.046 | 1791 | 0.901 | 0.905 |
|  | ReqSens 95% | 0.010 | 6269 | 0.950 | 0.655 |
| ^1^The method used to optimize thresholds based on the PresenceAbsence package in R. | | | | | |
| ^2^Sens=Spec finds the threshold where sensitivity equals specificity  ^3^MaxSensSpec finds the threshold that maximizes the sum of sensitivity and specificity. | | | | | |
| ^4^MaxKappa finds the threshold that maximizes the Kappa coefficient, a measure of the agreement between the predicted and observed values. | | | | | |
| ^5^MaxPCC finds the threshold that maximizes the Pearson correlation coefficient (PCC) between two vectors. | | | | | |
| ^6^ReqSens finds the threshold that meets the minimum sensitivity as specified by the user. | | | | | |

| **Table S6. Importance and ranks of ecological (*n* = 63) and taxonomic (*n* = 110) traits of mammal hosts for the host exposure model (based on molecular detection of viral genomes using PCR techniques) and the susceptible host model (based on virus isolation from hosts).** | | | | |
| --- | --- | --- | --- | --- |
|  | Importance | | Rank | |
| Feature | RT-PCR | Virus isolation | RT-PCR | Virus isolation |
| activity_crepuscular | 0.005704 | 0.006971 | 36 | 28 |
| activity_diurnal | 0.007585 | 0.007256 | 29 | 27 |
| activity_nocturnal | 0.00119 | 0.00296 | 58 | 48 |
| adult_body_length_mm | 0.009025 | 0.005299 | 27 | 36 |
| adult_mass_g | 0.021185 | 0.008059 | 13 | 26 |
| age_first_reproduction_d | 0.012332 | 0.006865 | 20 | 31 |
| altitude_breadth_m | 0.021688 | 0.020472 | 11 | 10 |
| biogeo_afrotropical | 0.006548 | 0.011104 | 34 | 20 |
| biogeo_antarctic | 0.001574 | 0 | 53 | 78 |
| biogeo_australasian | 0.002918 | 0.0016 | 45 | 52 |
| biogeo_indomalayan | 0.009716 | 0.006137 | 25 | 33 |
| biogeo_nearctic | 0.004778 | 0.004504 | 39 | 39 |
| biogeo_neotropical | 0.005228 | 0.006914 | 37 | 29 |
| biogeo_oceanian | 0.001817 | 0.000135 | 50 | 64 |
| biogeo_palearctic | 0.005833 | 0.011599 | 35 | 18 |
| brain_mass_g | 0.018855 | 0.008454 | 16 | 24 |
| cites | 0.293321 | 0.265897 | 1 | 1 |
| det_diet_breadth_n | 0.006916 | 0.019135 | 32 | 11 |
| det_fruit | 0.007795 | 0.00394 | 28 | 40 |
| det_inv | 0.006994 | 0.006873 | 31 | 30 |
| det_nect | 0.003881 | 0.011139 | 42 | 19 |
| det_plantother | 0.010818 | 0.023184 | 22 | 9 |
| det_scav | 0.002684 | 0.000536 | 46 | 57 |
| det_seed | 0.007293 | 0.016536 | 30 | 13 |
| det_vect | 0.000155 | 0.000232 | 65 | 63 |
| det_vend | 0.010875 | 0.017241 | 21 | 12 |
| det_vfish | 0.013596 | 0.000433 | 18 | 59 |
| det_vunk | 0.001415 | 0.00107 | 56 | 54 |
| disected_by_mountains | 0.013733 | 0.00975 | 17 | 21 |
| dispersal_km | 0.046592 | 0.13452 | 4 | 2 |
| dphy_invertebrate | 0.002943 | 0.004871 | 44 | 38 |
| dphy_plant | 0.00391 | 0.003294 | 41 | 44 |
| dphy_vertebrate | 0.010657 | 0.003597 | 23 | 43 |
| ed_equal | 0.038086 | 0.028703 | 5 | 8 |
| fam_ABROCOMIDAE | 0 | 0 | 86 | 79 |
| fam_AILURIDAE | 0 | 0 | 87 | 80 |
| fam_ANOMALURIDAE | 0 | 0 | 88 | 81 |
| fam_ANTILOCAPRIDAE | 0 | 0 | 89 | 82 |
| fam_AOTIDAE | 0 | 0 | 90 | 83 |
| fam_APLODONTIIDAE | 0 | 0 | 91 | 84 |
| fam_Archaeolemuridae | 0 | 0 | 92 | 85 |
| fam_ATELIDAE | 0.000002 | 0.000309 | 83 | 62 |
| fam_BALAENIDAE | 0 | 0 | 93 | 86 |
| fam_BALAENOPTERIDAE | 0 | 0 | 94 | 87 |
| fam_BATHYERGIDAE | 0 | 0 | 95 | 88 |
| fam_BOVIDAE | 0.000023 | 0.000015 | 75 | 75 |
| fam_BRADYPODIDAE | 0 | 0 | 96 | 89 |
| fam_CALLITRICHIDAE | 0.00154 | 0.003009 | 54 | 47 |
| fam_CALOMYSCIDAE | 0 | 0 | 97 | 90 |
| fam_CAMELIDAE | 0.000009 | 0 | 80 | 91 |
| fam_CANIDAE | 0.000367 | 0.000004 | 62 | 77 |
| fam_CAPROMYIDAE | 0 | 0 | 98 | 92 |
| fam_CASTORIDAE | 0 | 0 | 99 | 93 |
| fam_CAVIIDAE | 0 | 0 | 100 | 94 |
| fam_CEBIDAE | 0.000014 | 0 | 78 | 95 |
| fam_CERCOPITHECIDAE | 0.000026 | 0.00002 | 74 | 73 |
| fam_CERVIDAE | 0.000032 | 0 | 73 | 96 |
| fam_CHEIROGALEIDAE | 0 | 0 | 101 | 97 |
| fam_CHINCHILLIDAE | 0 | 0 | 102 | 98 |
| fam_CRICETIDAE | 0.000044 | 0.000038 | 70 | 71 |
| fam_CTENODACTYLIDAE | 0 | 0 | 103 | 99 |
| fam_CTENOMYIDAE | 0 | 0 | 104 | 100 |
| fam_CUNICULIDAE | 0 | 0 | 105 | 101 |
| fam_CYCLOPEDIDAE | 0 | 0 | 106 | 102 |
| fam_DASYPROCTIDAE | 0 | 0 | 107 | 103 |
| fam_DAUBENTONIIDAE | 0 | 0 | 108 | 104 |
| fam_DELPHINIDAE | 0.009537 | 0 | 26 | 105 |
| fam_DIATOMYIDAE | 0 | 0 | 109 | 106 |
| fam_DIDELPHIDAE | 0.000002 | 0.000039 | 84 | 70 |
| fam_DINOMYIDAE | 0 | 0 | 110 | 107 |
| fam_DIPODIDAE | 0 | 0 | 111 | 108 |
| fam_ECHIMYIDAE | 0 | 0 | 112 | 109 |
| fam_ELEPHANTIDAE | 0 | 0.000005 | 113 | 76 |
| fam_EQUIDAE | 0 | 0 | 114 | 110 |
| fam_ERETHIZONTIDAE | 0 | 0 | 115 | 111 |
| fam_ERINACEIDAE | 0 | 0 | 116 | 112 |
| fam_ESCHRICHTIIDAE | 0 | 0 | 117 | 113 |
| fam_EUPLERIDAE | 0 | 0 | 118 | 114 |
| fam_FELIDAE | 0.002525 | 0.0402 | 48 | 5 |
| fam_GALAGIDAE | 0 | 0 | 119 | 115 |
| fam_GEOMYIDAE | 0 | 0 | 120 | 116 |
| fam_GIRAFFIDAE | 0 | 0 | 121 | 117 |
| fam_GLIRIDAE | 0 | 0 | 122 | 118 |
| fam_HERPESTIDAE | 0.000016 | 0 | 76 | 119 |
| fam_HETEROCEPHALIDAE | 0 | 0 | 123 | 120 |
| fam_HETEROMYIDAE | 0 | 0 | 124 | 121 |
| fam_HIPPOPOTAMIDAE | 0 | 0 | 125 | 122 |
| fam_HOMINIDAE | 0 | 0 | 126 | 123 |
| fam_HYAENIDAE | 0 | 0 | 127 | 124 |
| fam_HYLOBATIDAE | 0 | 0 | 128 | 125 |
| fam_HYSTRICIDAE | 0 | 0 | 129 | 126 |
| fam_INDRIIDAE | 0 | 0 | 130 | 127 |
| fam_INIIDAE | 0 | 0 | 131 | 128 |
| fam_LEMURIDAE | 0 | 0 | 132 | 129 |
| fam_LEPILEMURIDAE | 0 | 0 | 133 | 130 |
| fam_LEPORIDAE | 0.000015 | 0.000108 | 77 | 67 |
| fam_LORISIDAE | 0 | 0 | 134 | 131 |
| fam_Mammutidae | 0 | 0 | 135 | 132 |
| fam_Megaladapidae | 0 | 0 | 136 | 133 |
| fam_MEGALONYCHIDAE | 0 | 0 | 137 | 134 |
| fam_MEPHITIDAE | 0.000009 | 0.000111 | 81 | 66 |
| fam_MONODONTIDAE | 0 | 0 | 138 | 135 |
| fam_MOSCHIDAE | 0 | 0 | 139 | 136 |
| fam_MURIDAE | 0.000062 | 0.003711 | 69 | 42 |
| fam_MUSTELIDAE | 6.2E-05 | 0 | 68 | 137 |
| fam_Mylodontidae | 0 | 0 | 140 | 138 |
| fam_MYOCASTORIDAE | 0 | 0 | 141 | 139 |
| fam_MYRMECOPHAGIDAE | 0 | 0 | 142 | 140 |
| fam_NANDINIIDAE | 0 | 0 | 143 | 141 |
| fam_NEOBALAENIDAE | 0 | 0 | 144 | 142 |
| fam_NESOMYIDAE | 0.000011 | 0 | 79 | 143 |
| fam_NESOPHONTIDAE | 0 | 0 | 145 | 144 |
| fam_Nothrotheriidae | 0 | 0 | 146 | 145 |
| fam_OCHOTONIDAE | 0 | 0 | 147 | 146 |
| fam_OCTODONTIDAE | 0 | 0 | 148 | 147 |
| fam_ODOBENIDAE | 0 | 0 | 149 | 148 |
| fam_OTARIIDAE | 0.000002 | 0 | 85 | 149 |
| fam_PALAEOPROPITHECIDAE | 0 | 0 | 150 | 150 |
| fam_PEDETIDAE | 0 | 0 | 151 | 151 |
| fam_PETROMURIDAE | 0 | 0 | 152 | 152 |
| fam_PHOCIDAE | 0.000039 | 0.00002 | 72 | 72 |
| fam_PHOCOENIDAE | 0 | 0 | 153 | 153 |
| fam_PHYSETERIDAE | 0 | 0 | 154 | 154 |
| fam_PITHECIIDAE | 0 | 0 | 155 | 155 |
| fam_PLATACANTHOMYIDAE | 0 | 0 | 156 | 156 |
| fam_PLATANISTIDAE | 0 | 0 | 157 | 157 |
| fam_PRIONODONTIDAE | 0 | 0 | 158 | 158 |
| fam_PROCYONIDAE | 0.000006 | 0.000078 | 82 | 69 |
| fam_PROLAGIDAE | 0 | 0 | 159 | 159 |
| fam_RHINOCEROTIDAE | 0 | 0.000083 | 160 | 68 |
| fam_SCIURIDAE | 0.000043 | 0.00013 | 71 | 65 |
| fam_SOLENODONTIDAE | 0 | 0 | 161 | 160 |
| fam_SORICIDAE | 0 | 0 | 162 | 161 |
| fam_SPALACIDAE | 0 | 0 | 163 | 162 |
| fam_SUIDAE | 0 | 0 | 164 | 163 |
| fam_TALPIDAE | 0 | 0 | 165 | 164 |
| fam_TAPIRIDAE | 0 | 0 | 166 | 165 |
| fam_TARSIIDAE | 0 | 0 | 167 | 166 |
| fam_TAYASSUIDAE | 0 | 0 | 168 | 167 |
| fam_THRYONOMYIDAE | 0 | 0 | 169 | 168 |
| fam_TRAGULIDAE | 0 | 0 | 170 | 169 |
| fam_URSIDAE | 0 | 0 | 171 | 170 |
| fam_VIVERRIDAE | 0 | 0 | 172 | 171 |
| fam_ZIPHIIDAE | 7.9E-05 | 0 | 67 | 172 |
| female_maturity_d | 0.023516 | 0.008181 | 10 | 25 |
| forager_arboreal | 0.00062 | 0.000392 | 61 | 60 |
| forager_ground | 0.000997 | 0.002776 | 59 | 49 |
| forager_marine | 0 | 0 | 173 | 173 |
| forager_scansorial | 0.000098 | 0.000939 | 66 | 55 |
| fossoriality | 0.005111 | 0.006483 | 38 | 32 |
| freshwater | 0.001339 | 0.002238 | 57 | 50 |
| generation_length_d | 0.010086 | 0.00853 | 24 | 23 |
| gestation_length_d | 0.027224 | 0.054042 | 9 | 3 |
| glaciation | 0.003447 | 0.000887 | 43 | 56 |
| habitat_breadth_n | 0.019823 | 0.034288 | 15 | 7 |
| hibernation_torpor | 0.000885 | 0.000441 | 60 | 58 |
| interbirth_interval_d | 0.051729 | 0.01199 | 3 | 17 |
| island_dwelling | 0.030725 | 0.005606 | 7 | 35 |
| island_end_isolated | 0.00213 | 0.003274 | 49 | 45 |
| island_end_lgbridge | 0.006838 | 0.004955 | 33 | 37 |
| island_end_mainland | 0.002676 | 0.003248 | 47 | 46 |
| island_end_marine | 0.001691 | 0.000016 | 52 | 74 |
| litter_size_n | 0.021415 | 0.038202 | 12 | 6 |
| litters_per_year_n | 0.02781 | 0.012233 | 8 | 16 |
| lower_elevation_m | 0.013417 | 0.009712 | 19 | 22 |
| marine | 0.000255 | 0.000351 | 64 | 61 |
| max_longevity_d | 0.051924 | 0.016424 | 2 | 14 |
| terrestrial_non.volant | 0.00029 | 0.003777 | 63 | 41 |
| trophic_carnivores | 0.001519 | 0.001408 | 55 | 53 |
| trophic_herbivores | 0.001795 | 0.002135 | 51 | 51 |
| trophic_omnivores | 0.004243 | 0.005821 | 40 | 34 |
| upper_elevation_m | 0.035447 | 0.042197 | 6 | 4 |
| weaning_age_d | 0.020865 | 0.0123 | 14 | 15 |

| **Table S7. Importance and rank of mammal traits (*N =* 173) and viral features (N = 10) for the link prediction model trained on host and viral traits.** Mammal traits include 63 ecological traits and 110 taxonomic traits. | | |
| --- | --- | --- |
| Feature | Importance | Rank |
| activity_crepuscular | 0.000163 | 80 |
| activity_diurnal | 0.000204 | 77 |
| activity_nocturnal | 0.000674 | 52 |
| adult_body_length_mm | 0.004499 | 20 |
| adult_mass_g | 0.004937 | 19 |
| age_first_reproduction_d | 0.005072 | 18 |
| altitude_breadth_m | 0.003835 | 23 |
| biogeo_afrotropical | 0.000387 | 66 |
| biogeo_antarctic | 0.000051 | 91 |
| biogeo_australasian | 0.000358 | 69 |
| biogeo_indomalayan | 0.001522 | 41 |
| biogeo_nearctic | 0.000286 | 71 |
| biogeo_neotropical | 0.000418 | 64 |
| biogeo_oceanian | 0.000267 | 73 |
| biogeo_palearctic | 0.000401 | 65 |
| brain_mass_g | 0.001691 | 40 |
| cites | 0.349742 | 1 |
| det_diet_breadth_n | 0.003384 | 27 |
| det_fruit | 0.000711 | 51 |
| det_inv | 0.000504 | 58 |
| det_nect | 0.001819 | 39 |
| det_plantother | 0.001928 | 36 |
| det_scav | 0.003768 | 24 |
| det_seed | 0.000284 | 72 |
| det_vect | 0.00048 | 59 |
| det_vend | 0.002261 | 33 |
| det_vfish | 0.003522 | 25 |
| det_vunk | 0.001519 | 42 |
| disected_by_mountains | 0.001254 | 44 |
| dispersal_km | 0.017614 | 9 |
| dphy_invertebrate | 0.0002 | 78 |
| dphy_plant | 0.000542 | 57 |
| dphy_vertebrate | 0.000823 | 49 |
| ed_equal | 0.00587 | 16 |
| fam_ABROCOMIDAE | 0.000002 | 106 |
| fam_AILURIDAE | 0.001834 | 37 |
| fam_ANOMALURIDAE | 0.000004 | 101 |
| fam_ANTILOCAPRIDAE | 0 | 110 |
| fam_AOTIDAE | 0 | 111 |
| fam_APLODONTIIDAE | 0 | 112 |
| fam_Archaeolemuridae | 0 | 113 |
| fam_ATELIDAE | 0.000088 | 87 |
| fam_BALAENIDAE | 0.002605 | 30 |
| fam_BALAENOPTERIDAE | 0 | 114 |
| fam_BATHYERGIDAE | 0 | 115 |
| fam_BOVIDAE | 0.000081 | 88 |
| fam_BRADYPODIDAE | 0 | 116 |
| fam_CALLITRICHIDAE | 0.000261 | 75 |
| fam_CALOMYSCIDAE | 0 | 117 |
| fam_CAMELIDAE | 0.003481 | 26 |
| fam_CANIDAE | 0.000479 | 60 |
| fam_CAPROMYIDAE | 0 | 118 |
| fam_CASTORIDAE | 0 | 119 |
| fam_CAVIIDAE | 0 | 120 |
| fam_CEBIDAE | 0.000979 | 46 |
| fam_CERCOPITHECIDAE | 0.000479 | 61 |
| fam_CERVIDAE | 0 | 121 |
| fam_CHEIROGALEIDAE | 1E-06 | 109 |
| fam_CHINCHILLIDAE | 0 | 122 |
| fam_CRICETIDAE | 0.000002 | 107 |
| fam_CTENODACTYLIDAE | 0 | 123 |
| fam_CTENOMYIDAE | 0 | 124 |
| fam_CUNICULIDAE | 0 | 125 |
| fam_CYCLOPEDIDAE | 0 | 126 |
| fam_DASYPROCTIDAE | 0 | 127 |
| fam_DAUBENTONIIDAE | 0 | 128 |
| fam_DELPHINIDAE | 0.001825 | 38 |
| fam_DIATOMYIDAE | 0 | 129 |
| fam_DIDELPHIDAE | 0.00004 | 93 |
| fam_DINOMYIDAE | 0 | 130 |
| fam_DIPODIDAE | 0 | 131 |
| fam_ECHIMYIDAE | 0 | 132 |
| fam_ELEPHANTIDAE | 0.000593 | 53 |
| fam_EQUIDAE | 0.000042 | 92 |
| fam_ERETHIZONTIDAE | 0 | 133 |
| fam_ERINACEIDAE | 0.000002 | 103 |
| fam_ESCHRICHTIIDAE | 0 | 134 |
| fam_EUPLERIDAE | 0 | 135 |
| fam_FELIDAE | 0.002674 | 29 |
| fam_GALAGIDAE | 0 | 136 |
| fam_GEOMYIDAE | 0 | 137 |
| fam_GIRAFFIDAE | 0.000587 | 55 |
| fam_GLIRIDAE | 0.000002 | 105 |
| fam_HERPESTIDAE | 0 | 138 |
| fam_HETEROCEPHALIDAE | 0 | 139 |
| fam_HETEROMYIDAE | 0 | 140 |
| fam_HIPPOPOTAMIDAE | 0 | 141 |
| fam_HOMINIDAE | 0.000322 | 70 |
| fam_HYAENIDAE | 0 | 142 |
| fam_HYLOBATIDAE | 0.000067 | 89 |
| fam_HYSTRICIDAE | 0 | 143 |
| fam_INDRIIDAE | 0 | 144 |
| fam_INIIDAE | 0.000025 | 95 |
| fam_LEMURIDAE | 0 | 145 |
| fam_LEPILEMURIDAE | 0 | 146 |
| fam_LEPORIDAE | 0.000005 | 100 |
| fam_LORISIDAE | 0 | 147 |
| fam_Mammutidae | 0 | 148 |
| fam_Megaladapidae | 0 | 149 |
| fam_MEGALONYCHIDAE | 0 | 150 |
| fam_MEPHITIDAE | 0.000115 | 83 |
| fam_MONODONTIDAE | 0 | 151 |
| fam_MOSCHIDAE | 0 | 152 |
| fam_MURIDAE | 0.000368 | 68 |
| fam_MUSTELIDAE | 0.000002 | 104 |
| fam_Mylodontidae | 0 | 153 |
| fam_MYOCASTORIDAE | 0 | 154 |
| fam_MYRMECOPHAGIDAE | 0.002143 | 34 |
| fam_NANDINIIDAE | 0 | 155 |
| fam_NEOBALAENIDAE | 0.00001 | 97 |
| fam_NESOMYIDAE | 0 | 156 |
| fam_NESOPHONTIDAE | 0 | 157 |
| fam_Nothrotheriidae | 0 | 158 |
| fam_OCHOTONIDAE | 0 | 159 |
| fam_OCTODONTIDAE | 0 | 160 |
| fam_ODOBENIDAE | 0 | 161 |
| fam_OTARIIDAE | 0.000003 | 102 |
| fam_PALAEOPROPITHECIDAE | 0 | 162 |
| fam_PEDETIDAE | 0 | 163 |
| fam_PETROMURIDAE | 0 | 164 |
| fam_PHOCIDAE | 0 | 165 |
| fam_PHOCOENIDAE | 0.00004 | 94 |
| fam_PHYSETERIDAE | 0 | 166 |
| fam_PITHECIIDAE | 0 | 167 |
| fam_PLATACANTHOMYIDAE | 0 | 168 |
| fam_PLATANISTIDAE | 0 | 169 |
| fam_PRIONODONTIDAE | 0 | 170 |
| fam_PROCYONIDAE | 0.000478 | 62 |
| fam_PROLAGIDAE | 0 | 171 |
| fam_RHINOCEROTIDAE | 0.000157 | 81 |
| fam_SCIURIDAE | 0.000248 | 76 |
| fam_SOLENODONTIDAE | 0 | 172 |
| fam_SORICIDAE | 0.000001 | 108 |
| fam_SPALACIDAE | 0 | 173 |
| fam_SUIDAE | 0 | 174 |
| fam_TALPIDAE | 0 | 175 |
| fam_TAPIRIDAE | 0.000947 | 47 |
| fam_TARSIIDAE | 0 | 176 |
| fam_TAYASSUIDAE | 0 | 177 |
| fam_THRYONOMYIDAE | 0 | 178 |
| fam_TRAGULIDAE | 0 | 179 |
| fam_URSIDAE | 0 | 180 |
| fam_VIVERRIDAE | 0 | 181 |
| fam_ZIPHIIDAE | 0 | 182 |
| female_maturity_d | 0.002088 | 35 |
| forager_arboreal | 0.000589 | 54 |
| forager_ground | 0.000009 | 99 |
| forager_marine | 0 | 183 |
| forager_scansorial | 0.000106 | 85 |
| fossoriality | 0.000752 | 50 |
| freshwater | 0.000014 | 96 |
| generation_length_d | 0.00138 | 43 |
| gestation_length_d | 0.007116 | 13 |
| glaciation | 0.000056 | 90 |
| habitat_breadth_n | 0.001086 | 45 |
| hibernation_torpor | 0.000907 | 48 |
| interbirth_interval_d | 0.002734 | 28 |
| island_dwelling | 0.041044 | 6 |
| island_end_isolated | 0.000547 | 56 |
| island_end_lgbridge | 0.000267 | 74 |
| island_end_mainland | 0.000175 | 79 |
| island_end_marine | 0.00013 | 82 |
| litter_size_n | 0.006431 | 14 |
| litters_per_year_n | 0.005588 | 17 |
| lower_elevation_m | 0.002437 | 32 |
| marine | 0.00001 | 98 |
| max_longevity_d | 0.002439 | 31 |
| PC1 | 0.117091 | 3 |
| PC10 | 0.040512 | 7 |
| PC2 | 0.012221 | 11 |
| PC3 | 0.053417 | 4 |
| PC4 | 0.147444 | 2 |
| PC5 | 0.00447 | 21 |
| PC6 | 0.004406 | 22 |
| PC7 | 0.023601 | 8 |
| PC8 | 0.013748 | 10 |
| PC9 | 0.05098 | 5 |
| terrestrial_non.volant | 0.0001 | 86 |
| trophic_carnivores | 0.000458 | 63 |
| trophic_herbivores | 0.000107 | 84 |
| trophic_omnivores | 0.000379 | 67 |
| upper_elevation_m | 0.005965 | 15 |
| weaning_age_d | 0.00717 | 12 |

Dataset S1. Host-virus associations for link prediction based on Orthopoxvirus genomes extracted from NCBI.

Dataset S2. Predictive function of associated proteins for genes ranked by their positive and negative loading contribution to principal components 1, 4, 3 and 9. Genes with loading values that were greater than the mean plus 1.5 times the standard deviation (SD) and less than the mean minus the SD were included for those with positive loadings separate from those with negative loadings.

**SI References**

1.	Upham NS, Esselstyn JA, Jetz W. Inferring the mammal tree: Species-level sets of phylogenies for questions in ecology, evolution, and conservation. PLOS Biology. 2019;17: e3000494. doi:10.1371/journal.pbio.3000494

2.	Katoh K, Standley DM. MAFFT Multiple Sequence Alignment Software Version 7: Improvements in Performance and Usability. Molecular Biology and Evolution. 2013;30: 772–780. doi:10.1093/molbev/mst010

3.	Katoh K, Misawa K, Kuma K, Miyata T. MAFFT: a novel method for rapid multiple sequence alignment based on fast Fourier transform. Nucleic Acids Research. 2002;30: 3059–3066. doi:10.1093/nar/gkf436

4.	The IUCN Red List of Threatened Species. In: IUCN Red List of Threatened Species [Internet]. [cited 14 Jul 2023]. Available: https://www.iucnredlist.org/en

5.	Home - Taxonomy - NCBI. [cited 14 Jul 2023]. Available: https://www.ncbi.nlm.nih.gov/taxonomy

6.	Pardiñas UFJ, Lessa G, Teta P, Salazar-Bravo J, Câmara EMVC. A new genus of sigmodontine rodent from eastern Brazil and the origin of the tribe Phyllotini. Journal of Mammalogy. 2014;95: 201–215. doi:10.1644/13-MAMM-A-208
